# Learning Interpretable Switching Dynamics in Shared Neural-Behavioral Latent Space

**DOI:** 10.64898/2026.10.02.756323

**Authors:** Yongxu Zhang, Josue Ortega Caro, Rachel Oren, Michael James Higley, Jessica Cardin, Anne Churchland, Shreya Saxena

**Affiliations:** Yale University; UCLA

## Abstract

Modern recording technologies enable simultaneous measurement of high-dimensional neural activity and rich behavioral variables, creating both an opportunity and a modeling challenge: identifying the shared latent structure that mediates the brain-behavior relationship and the underlying dynamics through which it evolves across behavioral states. Existing approaches focus on different parts of this challenge: shared representation learning methods identify neural-behavioral subspaces but typically do not model dynamical structure, while dynamical models capture temporal variation and discrete state transitions in neural activity, but do not explicitly disentangle shared neural-behavioral structure from modality-specific variability. We introduce Switching Shared Latent Dynamics (SSLD), a unified framework that learns a latent representation shared between neural activity and behavior and endows this shared space with nonlinear switching recurrent dynamics to reflect the hypothesis that behaviorally relevant neural representations exhibit structured dynamical changes across behavioral states. Complementary private latent variables capture modality-specific variability, providing a window into neural activity that does not directly interact with behavior. We evaluate SSLD on a simulated dataset and four diverse experimental datasets spanning species, brain regions, and recording modalities: monkey motor and premotor cortex during reaching, somatosensory cortex during a bump task, widefield calcium imaging across mouse dorsal cortex during (a) self-initiated decision-making and (b) spontaneous movements. Across all four datasets, SSLD accurately reconstructs neural and behavioral signals, recovers discrete states in the dynamics that align with experimentally defined behavioral epochs, and isolates behaviorally-relevant neural information in the shared latent while preserving private neural variability. Ablations confirm that shared representation learning, behavioral supervision, and switching dynamics each contribute to performance. SSLD offers an interpretable approach to modeling shared neural-behavioral dynamics that complements ongoing efforts toward foundation models for neuroscience.

## 1 Introduction

Understanding how neural population activity gives rise to behavior is a central goal of systems neuroscience. This goal has been advanced by recent recording technologies that capture neural population activity together with detailed behavioral measurements. Such technologies offer an opportunity to uncover the latent structure linking brain activity to behavior, but they also introduce a major modeling challenge: neural activity contains both behaviorally relevant signals and substantial variability unrelated to the measured behavior. Therefore, a useful model should not only accurately represent neural and behavioral data, but also disentangle shared latent factors from neural activity that does not influence ongoing behavior.

One line of work has tackled this problem by modeling neural activity on its own. Latent dynamical systems infer continuous trajectories underlying neural population activity [Linderman et al., 2017, Pandarinath et al., 2018, Dong et al., 2020, Glaser et al., 2020, Durstewitz et al., 2023, Karniol-Tambour et al., 2024, Zhang and Saxena, 2024]. Such models can reveal rich structure in neural dynamics, but without behavioral grounding, the resulting latent dynamics are not guaranteed to align with the behavioral states that animals are actually moving through. A complementary line of work has incorporated behavioral variables directly into representation learning [Schneider et al., 2023, Sani et al., 2021, 2024, Yi et al., 2025]. However, they typically treat behavior as an additional observation to be predicted or aligned at the level of activity, without modeling how the shared neural-behavioral representation evolves in time. As a result, behavioral alignment is imposed on the static structure of the shared subspace, but not on its dynamics - yet the feature that makes neural activity behaviorally meaningful is precisely how it changes across behavioral epochs, cognitive states, and task phases.

Here, to address this important gap, we introduce a Switching Shared Latent Dynamics (SSLD). SSLD learns a latent representation shared between neural activity and behavior, while endowing this shared latent space with switching recurrent dynamics. This design is motivated by the hypothesis that if a latent neural representation is behaviorally relevant, then its dynamics should change in a structured way across behavioral states. By learning switching dynamics specifically in the shared latent subspace, the model focuses its discrete state inference on the component of neural activity most directly linked to behavior, rather than on arbitrary neural variability.

We evaluate SSLD on four datasets covering different species, brain regions, behavioral paradigms, and neural recording modalities. Our experiments demonstrate that: **(1)** SSLD accurately identifies the shared information between neural activity and behavior; **(2)** the discrete switches inferred within the shared latent space align significantly better with ground-truth behavioral states than switches operating on neural activity alone; **(3)** the combination of switching dynamics and shared-private disentanglement yields substantially more interpretable latent representations, with each regime capturing a distinct phase of neural-behavioral coupling; and **(4)** the model’s modular structure – with explicit switching dynamics – represents a meaningful step toward more interpretable and scalable models of brain-behavior relationships. Our work contributes to a broader effort toward building *foundation models for neuroscience* – flexible, multi-modal architectures that can be applied across diverse neural datasets, brain regions, and behavioral contexts [Ye et al., 2023, Azabou et al., 2023, Zhang et al., 2024, Azabou et al., 2025, Ryoo et al., 2025]. While recent foundation models for neural data have focused primarily on scaling across sessions and subjects, they have largely treated dynamics as stationary. By demonstrating that switching dynamics in a shared latent space provide a principled and interpretable bridge between neural activity and behavior, SSLD offers a complementary architectural motif that we believe will be essential for the next generation of neuroscience foundation models.

## 2 Related Works

### Multi-modal models

Several recent methods incorporate behavior into neural representation learning. CEBRA learns nonlinear latent embeddings from joint behavioral and neural data, enabling consistent behaviorally informed representations across tasks and species [Schneider et al., 2023], while PSID and DPAD separate behaviorally relevant neural dynamics from other neural variability [Sani et al., 2021, 2024]. Multi-modal Gaussian process variational autoencoders (MM-GPVAE) further model shared neural-behavioral structure using flexible, nonparametric latent dynamics [Gondur et al., 2024]. Shared-AE learns latent features shared between neural activity and behavior while separating modality-private features [Yi et al., 2025], providing a framework for isolating neural-behavioral shared representations. These approaches highlight the importance of isolating neural dynamics predictive of behavior, but they generally do not explicitly model discrete switching structure in the learned behaviorally relevant latent space.

### Switching dynamical systems

In parallel, switching dynamical systems model time series through transitions among discrete dynamical regimes, including SLDS and recurrent SLDS variants [Lin-derman et al., 2017]. Switching nonlinear dynamics such as SNLDS, MRSDS and SRNNs, extends this idea by using nonlinear neural networks whose weights switch over time, enabling nonlinear modeling of switching neural dynamics [Dong et al., 2020, Karniol-Tambour et al., 2024, Zhang and Saxena, 2024]. However, these switching nonlinear dynamical systems infer switches directly from neural activity without explicitly tying them to behaviorally shared latent structure.

### Foundation models in neuroscience

Finally, recent work has begun to develop foundation models for neuroscience: large, pretrained architectures designed to generalize across sessions, subjects, brain regions, and tasks. POYO introduced a unified, scalable framework for neural population decoding [Azabou et al., 2023], later extended to multi-session, multi-task decoding across distinct cell types and brain regions [Azabou et al., 2025], and further advanced through hybrid state-space modeling [Ryoo et al., 2025]. Related efforts include BrainLM, which applies masked autoencoding to large-scale fMRI recordings [Caro et al., 2024]; “universal translator” models that learn transferable representations of spike trains at single-cell, single-spike resolution [Zhang et al., 2024]; a unified framework for neural encoding and decoding at scale [Zhang et al., 2025]; and the Population Transformer, which learns flexible population-level representations across heterogeneous channel configurations [Chau et al., 2025]. While these models have made significant progress in scaling across sessions and subjects, they have largely treated neural dynamics as stationary and offer limited interpretability of the underlying neural dynamics.

## 3 Methods

SSLD learns a shared neural-behavioral latent representation while placing nonlinear switching recurrent dynamics within it (Fig. 1). This allows the inferred states to capture interpretable dynamical transitions associated with the component of neural activity most aligned with behavior. Moreover, the separate shared vs. private spaces enable the model to disentangle shared neural-behavioral representations from modality-specific variability.

**Figure 1:**
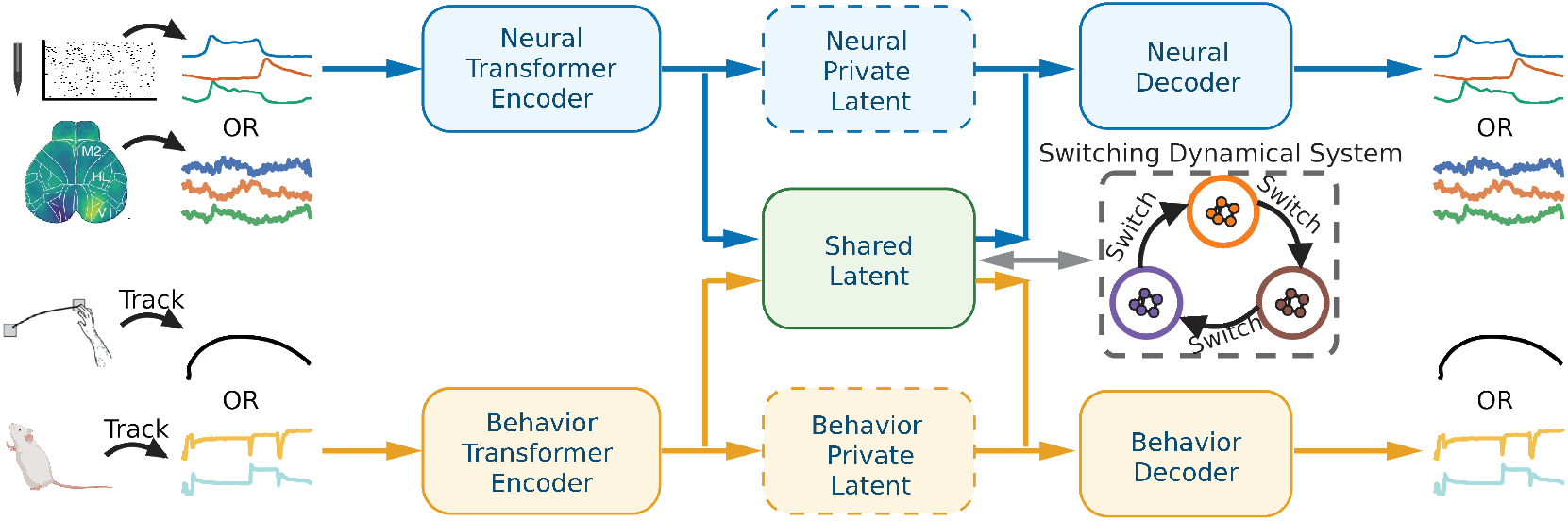
Overview of Switching Shared Latent Dynamics (SSLD). Neural activity and behavior are encoded by separate Transformer-based encoders into a shared latent representation and modality-specific private latent variables. The shared latent is modeled with switching dynamics to capture behaviorally-relevant discrete dynamical regimes. The shared and private modality-specific latent representations are then used to reconstruct the corresponding neural and behavioral signals.

Let 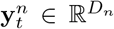 and 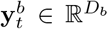 denote the neural activity and behavioral observations at time *t*, respectively. Our model decomposes the information in each modality into three latent components: **(1)** A *shared latent* 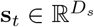 that captures information common to both neural activity and behavior and evolves according to switching nonlinear dynamics; **(2)** A *private neural latent* 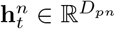 that captures neural variability not explained by behavior; **(3)** A *private behavioral latent* 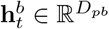 that captures behavioral variability not explained by the recorded neural activity. The shared latent is equipped with switching recurrent dynamics, enabling it to capture discrete regime changes in joint neural-behavioral activity.

### 3.1 Generative Model

#### Switching dynamics on the shared latent

The shared latent **s**_*t*_ evolves according to a Switching Recurrent Neural Network (SRNN) [Zhang and Saxena, 2024]. At each *t*, a discrete switching state **z**_*t*_ *∈ {*1, …, *K}* selects one of *K* distinct nonlinear dynamical regimes. The generative process is:

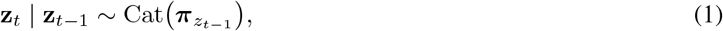

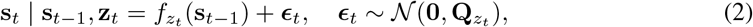

where 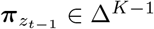 is the **z**_*t−*1_-th row of a *K × K* Markov transition matrix, *f*_*z*_ (*·*) is the RNN transition function with regime-specific weights 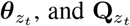 is the process noise covariance for regime **z**_*t*_. For the initial time step, **z**_0_ *∼* Cat(***π***_0_). Here, the dynamical model is flexible and can be adapted under different assumptions, e.g., linear, nonlinear, switching, or non-switching dynamics.

#### Inference networks (encoder)

We build two inference networks for encoding neural and behavior latents respectively:

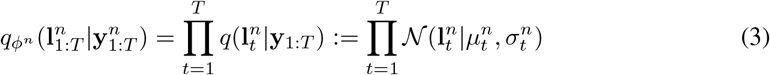

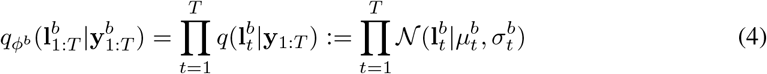

where *ϕ*^*n*^ and *ϕ*^*b*^ are parameters of inference networks for neural latent and behavior latent respectively, (*µ*^*n*^, *σ*^*n*^) and (*µ*^*b*^, *σ*^*b*^) are the means and variances of Gaussian distributions for neural latent and behavior latent respectively. Here, we use the reparameterization trick to infer the continuous hidden states [Kingma and Welling, 2013]:

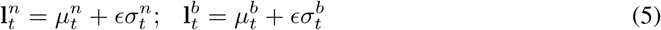

where *ϵ ∈ N* (0, 1) is a Gaussian noise parameter. The *µ*^*n*^ and *µ*^*b*^ are learned via training neural networks [Kingma and Welling, 2013]. In this work, we leverage Transformer-based encoders to generate them. We have

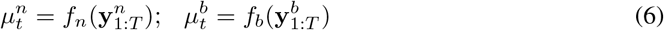

where *f*_*n*_ and *f*_*b*_ are Transformer encoders of neural and behavioral activity, respectively. The encoder architecture is flexible and can be adapted to other neural network models. In this paper, we report results using Transformer encoders; replacing them with multilayer perceptron (MLP) encoders yielded comparable performance.

#### Shared latents and private latents

The shared latent is extracted from encoded neural and behavioral latents together. The private latents are extracted deterministically by modality-specific encoders [Gondur et al., 2024, Yi et al., 2025]:

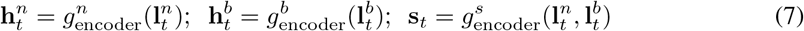

where 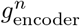 and 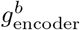 are linear networks parameterizing the private encoders for each modality, and 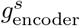 is a linear network parameterizing the shared encoder.

#### Observation model

Each modality is reconstructed from the concatenation of the shared latent and its corresponding private latent:

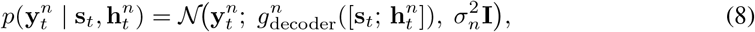

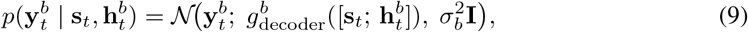

where 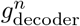 and 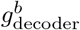 are decoder networks, and [ *·* ; *·* ] denotes concatenation.

### 3.2 Variational Inference

The shared latent **s**_1:*T*_ and switching states **z**_1:*T*_ are inferred via amortized variational inference. We introduce a factored variational posterior:

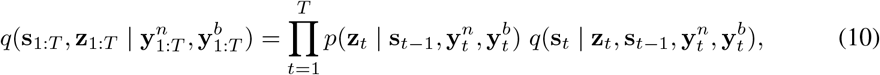

where *q*(**s** |**y**^*n*^, **y**^*b*^) is the posterior of shared latent given the observation. *p*(**z**| **s, y**^*n*^, **y**^*b*^) is the Bayesian posterior of discrete switching states given the observations. Note that unlike SRNN [Zhang and Saxena, 2024], which conditions only on neural observations, our inference network conditions on *both* modalities to infer the shared latent.

### 3.3 Loss Function

The total loss consists of two components: (1) a variational (ℒ_v_) for the switching shared dynamics along with the generative models for neural activity and behavior; (2) Cauchy-Schwarz (CS) divergence terms (*ℒ*_CS_) for shared-private disentanglement:

#### Variational ELBO loss (switching shared dynamics)

The evidence lower bound on log 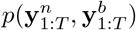 is :

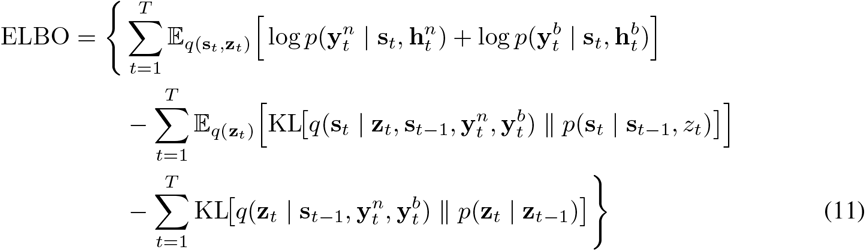

The first term encourages accurate reconstruction of both neural activity and behavior. The second term regularizes the shared latent dynamics to follow the switching RNN prior. The third term regularizes the inferred switching states to be consistent with the discrete behavioral states prior. We negate the ELBO so that the loss is minimized.

To encourage the inferred switches to capture behaviorally relevant structure, we use behavioral states as a prior over the switching states during training. Specifically, let 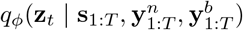 denote the inferred posterior distribution over the switching state **z**_*t*_, and let 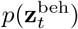 denote the behavioral-state prior at time *t*. We encourage the inferred switching distribution to align with this behavioral prior by minimizing the cross-entropy loss:

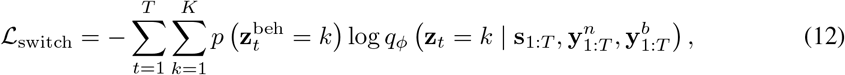

where *K* is the number of switching states. This loss encourages the model to infer switching states that are consistent with behaviorally defined state structure during training. **At test time, the model infers switching states directly from the data without access to the behavioral-state prior**. The overall variational loss *ℒ*_v_ is as following:

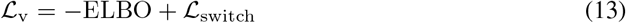

#### Disentanglement between latent representations

We encourage the private latents to capture information distinct from the shared latent. Since there are multiple ways to achieve this, we provide three options: *maximizing* the Cauchy–Schwarz (CS) divergence between the latent distributions [Yi et al., 2025], *maximizing* the orthogonality between the latent representations [Bousmalis et al., 2016], and *minimizing* the total correlation between the latents [Lee and Pavlovic, 2020]. The chosen option is applied between each private latent and the shared latent, as well as between the two private latents:

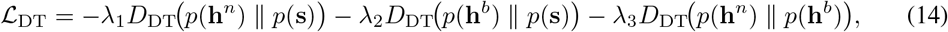

where *λ*_1_ *>* 0, *λ*_2_ *>* 0, and *λ*_3_ *>* 0 are hyperparameters controlling the strength of disentanglement regularization, and *D*_DT_ denotes the chosen disentanglement measure. Because ℒ_DT_ is minimized, each option is defined so that *D*_DT_ increases as the two latents become more disentangled. The first option is the CS divergence:

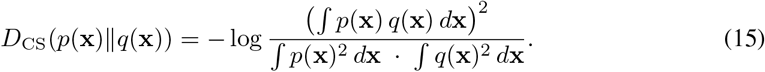

Since *D*_CS_*≥* 0 and equals zero when *p* = *q*, minimizing ℒ_CS_ (i.e., maximizing *D*_CS_) encourages the private latent distributions to diverge from the shared latent distribution, promoting disentanglement. In practice, we estimate *D*_CS_ using kernel density estimation with Gaussian kernels on mini-batch samples, following [Yi et al., 2025, Santana et al., 2016]. Note that unlike Shared-AE [Yi et al., 2025], which focuses on separating shared and private latent subspaces, our model emphasizes capturing temporal dynamics and their discrete switching structure.

The second option is orthogonality. Given a minibatch of *M* inputs, we stack the corresponding samples from *p*(**x**) and *q*(**y**) row-wise into matrices 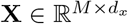 and 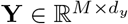, and apply a soft orthogonality constraint:

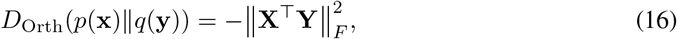

where *∥ · ∥*_*F*_ is the Frobenius norm. This term reaches its maximum of zero when every dimension of **x** is orthogonal to every dimension of **y** across the minibatch.

The third option is total correlation. Following Lee and Pavlovic [2020], who penalize total correlation to make private and shared factors statistically independent, we use the negative total correlation between the two latents:

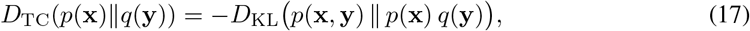

where *p*(**x, y**) is the joint distribution of the two latents, with marginals *p*(**x**) and *q*(**y**). This term reaches its maximum of zero when **x** and **y** are independent. We estimate it with minibatch-weighted sampling, as in Lee and Pavlovic [2020].

#### Full objective

The complete training objective, which is minimized over all encoder, decoder, and dynamics parameters, is:

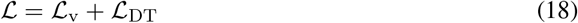

In the main text, we mainly report results using the CS divergence; comparisons across alternative objectives are provided in Table K.1. The model is trained end-to-end using the Adam optimizer.

#### Reproducibility statement

Our implementation is based on Pytorch 2.8.0 and we train our models using NVIDIA A100 GPUs. All experimental datasets are publicly accessible (see Appendix B). We provide the analysis and the results in this paper; the original code for the entire SSLD framework on Pytorch has been made public https://github.com/saxenalab-neuro/SSLD

## 4 Result

We detail the results of SSLD on a simulated dataset and four experimental datasets ranging from monkey electrophysiology (EPhys) to mouse widefield calcium imaging (WFCI). Unless specified, **all quantitative results are reported on held-out test sets**.

### 4.1 Simulated Data

We first evaluate SSLD on simulated data. The details of simulated data is shown in Appendix A. As shown in Fig. M.1, SSLD accurately reconstructs both neural activity and behavior, infers correct switching states, and recovers the ground-truth shared latent.

### 4.2 Monkey Reaching Task with EPhys from M1 and PMd

We apply SSLD on the firing rates of single units recorded from monkey motor and premotor cortex (*D*_*n*_ = 180), while the monkey performs a reaching task in 27 reaching conditions given targets at different locations. Neural activity was averaged across trials under the same condition. The behavior data is from hand movement, hand velocity, and muscle activity (*D*_*b*_ = 13). More details can be found in Appendix B and [Churchland et al., 2012]. We use 5-fold cross validation.

SSLD accurately reconstructed both neural population activity and behavioral trajectories, as illustrated by representative examples in Fig. 2A and Fig. 2B and quantified by mean squared error in Table G.1.. We next examined how behavioral information was distributed across the learned representations (Appendix E). We used a linear decoder to assess how directly the latent representations encode behavior. Behavioral information was strongly represented in the shared latent and private behavioral latent, but was largely absent from the private neural latent (Fig. 2C), indicating that SSLD separates behaviorally relevant shared structure from neural-specific variability. We report decoding performance using an long-short term memory (LSTM) decoder in Table H.1. We further tested whether the shared latent captured the global geometry of the task by comparing its condition-wise correlation structure with a behavioral condition-similarity matrix, defined by the inverse Euclidean distance between source-to-target reaching paths (Fig. 2D). These structures were highly aligned, with a correlation coefficient of *r* = 0.942, showing that conditions with similar task geometry produced more similar shared latents (details in Appendix F and Fig. M.6). The shared latent showed stronger correspondence with task geometry and behavioral condition similarity, than raw neural activity or private latents (Fig. 2E).

**Figure 2:**
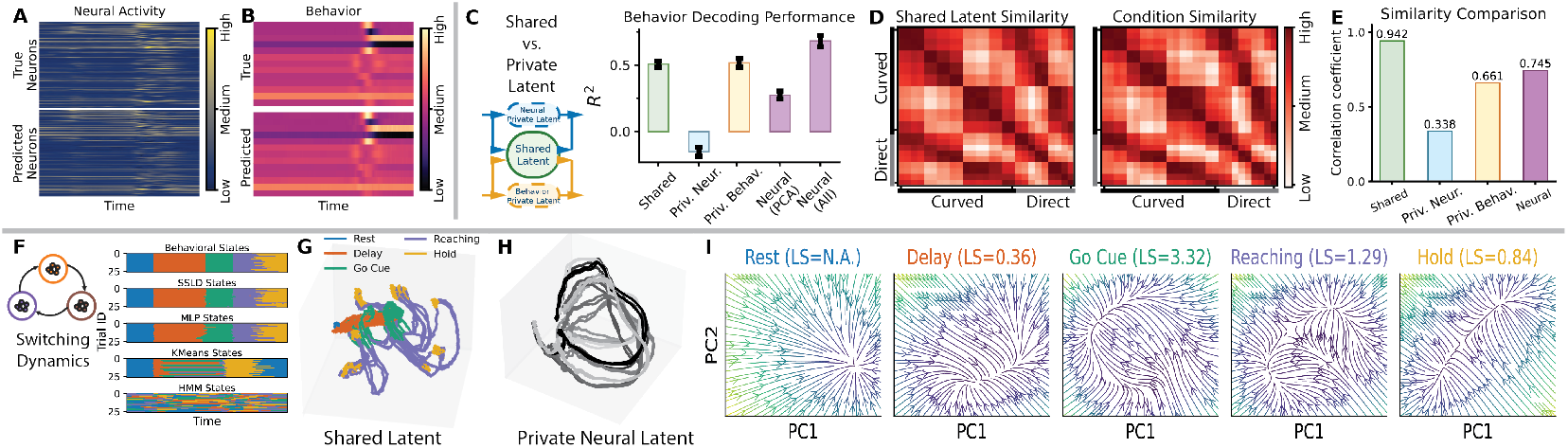
Reaching: Representative reconstructions of (A) neural activity and (B) behavior. *Shared vs. private latent*: (C) Behavioral information is mainly encoded in the shared and private behavioral latents, as assessed using a linear decoder (PN: private neural latent; PB: private behavior latent). Behavior decoding performance using latent and LSTM decoder is shown in Table H.1 (D) The shared-latent similarity matrix and condition similarity across the 27 reaching conditions. (E) Linear regression coefficients relating condition similarity to similarity in the shared latent, private neural latent, private behavioral latent, and raw neural activity, respectively (see Appendix F). *Switching dynamics*: (F) SSLD inferred discrete states align with task-defined behavioral epochs. Baselines include a supervised MLP and unsupervised methods (*k*-means and HMM). (G) Shared latent trajectories show structured organization across conditions. (H) Private neural latent trajectories show large rotational signals across conditions. (I) Flow fields of the RNN in distinct behavioral states, with line-attractor scores (LS) indicated in the titles, reveal state-dependent latent dynamics.

We then compared the discrete dynamical states inferred by SSLD with task-defined behavioral states (Fig. 2F). The inferred states closely aligned with behavioral epochs, and SSLD outperformed both unsupervised and supervised baselines (Table 1). Unsupervised baselines included *k*-means, hidden Markov models (HMMs), switching linear dynamical systems (SLDS), recurrent SLDS (rSLDS), and an SSLD variant without behavioral labels as a prior. Supervised baselines included a multilayer perceptron (MLP), an LSTM network, and a supervised SRNN. We used a shared latent dimensionality of *D*_*s*_ = 8 for the main analysis (see Appendix C) and further evaluated robustness across different values of *D*_*s*_ in Appendix Fig. M.2. Moreover, visualization of the shared latent trajectories in 3-d revealed clear trial-level structure in the inferred neural-behavioral representation (Fig. 2G). The private neural latent (Fig. 2H) captures condition-invariant structure in the neural activity. The private neural latent is highly correlated and rarely related to condition similarity, instead showing large rotational signals, which is consistent with prior findings [Kaufman et al., 2016] showing that a substantial component of population dynamics reflects condition-independent neural processes.

**Table 1:**
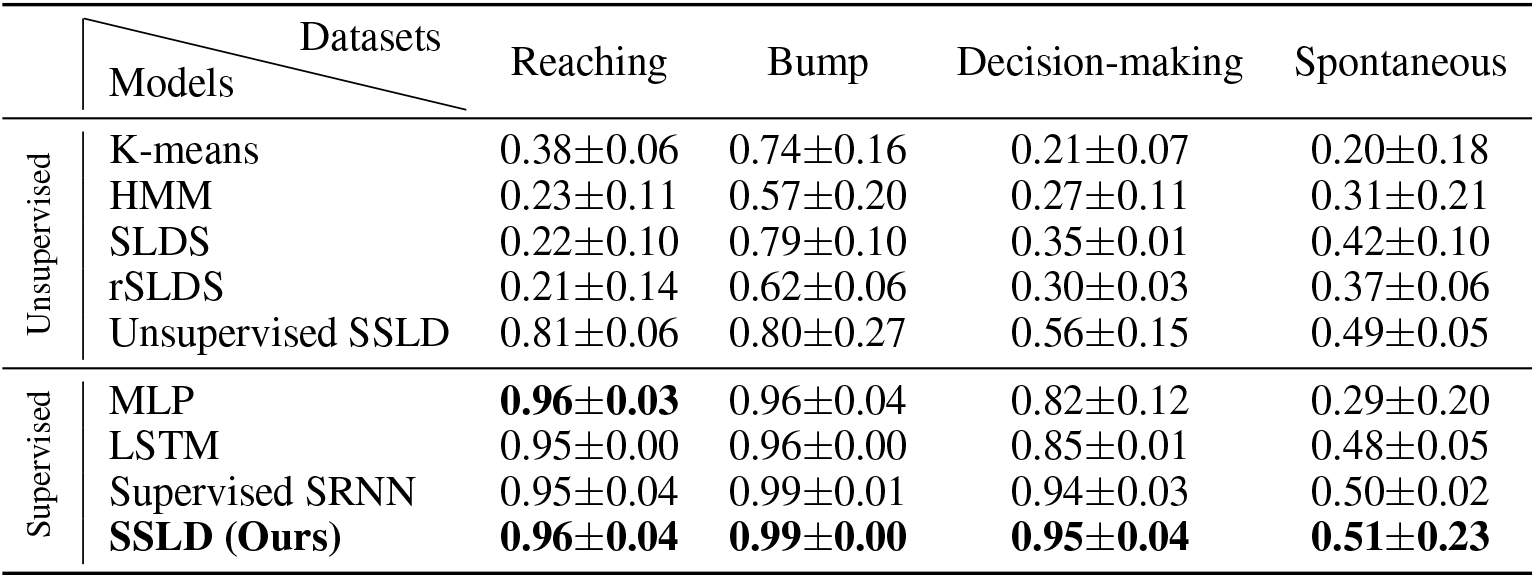
State Alignment Accuracy across Datasets (mean across trials*±*standard deviation)

|  | Datasets<br>Models | Reaching | Bump | Decision-making | Spontaneous |
|---|---|---|---|---|---|
| Unsupervised | K-means | 0.38±0.06 | 0.74±0.16 | 0.21±0.07 | 0.20±0.18 |
|  | HMM | 0.23±0.11 | 0.57±0.20 | 0.27±0.11 | 0.31±0.21 |
|  | SLDS | 0.22±0.10 | 0.79±0.10 | 0.35±0.01 | 0.42±0.10 |
|  | rSLDS | 0.21±0.14 | 0.62±0.06 | 0.30±0.03 | 0.37±0.06 |
|  | Unsupervised SSLD | 0.81±0.06 | 0.80±0.27 | 0.56±0.15 | 0.49±0.05 |
| Supervised | MLP | **0.96±0.03** | 0.96±0.04 | 0.82±0.12 | 0.29±0.20 |
|  | LSTM | 0.95±0.00 | 0.96±0.00 | 0.85±0.01 | 0.48±0.05 |
|  | Supervised SRNN | 0.95±0.04 | 0.99±0.01 | 0.94±0.03 | 0.50±0.02 |
|  | **SSLD (Ours)** | **0.96±0.04** | **0.99±0.00** | **0.95±0.04** | **0.51±0.23** |

To characterize the dynamics learned by SSLD, we visualized the flow field of the shared latent state *s* within each inferred discrete state (Fig. 2I). The flow fields differed across states, indicating distinct state-dependent regimes in the shared neural-behavioral latent space. We further quantified these differences using the line-attractor score (LS) of the learned RNN dynamics [Maheswaranathan et al., 2019, Vinograd et al., 2024]. The score increased after target onset (between ‘rest’ and ‘delay’), exceeded 1 following the ‘go cue’, and decreased after reaching, suggesting a transition from preparatory dynamics to a stable movement-execution regime and then back toward a more flexible baseline state.

Finally, we performed three ablations to assess the roles of the shared representation, behavioral observations, and switching dynamics. First, we applied Canonical Correlation Analysis (CCA) to neural activity and behavior after reducing both modalities to the shared-latent dimensionality (Table 2: Corr). Although CCA achieved high training correlation, its test performance dropped substantially, suggesting that we need nonlinear models to fully capture the neural-behavioral relationship. Second, removing the behavioral encoder and decoder reduced behavioral decoding from the learned latent representation (Table 2: *R*^2^), indicating that training the model with behavioral observations are important for capturing shared structure - oftentimes, the neural activity is dominated by recurrent activity that is not all relevant for ongoing behavior. Third, removing the switching recurrent dynamics and clustering the resulting latent states with *k*-means led to lower state-alignment accuracy than SSLD (Table 2: Acc), supporting the contribution of explicit switching dynamics during end-to-end model training.

**Table 2:**
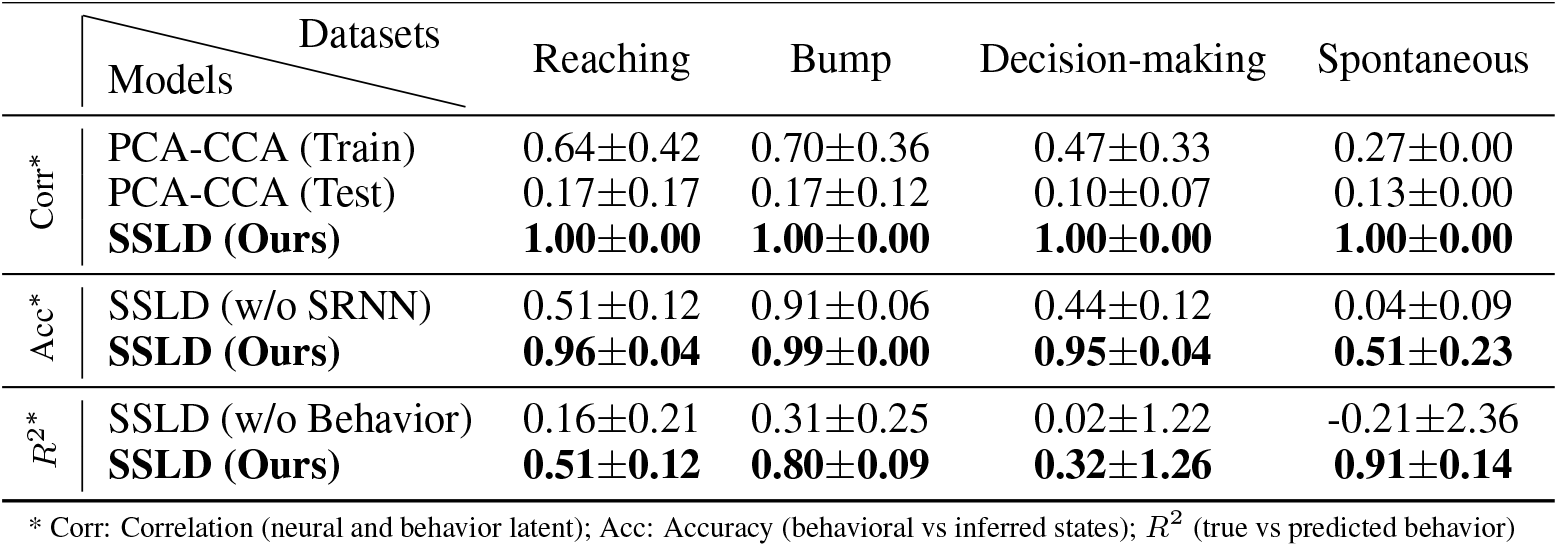
Ablation Test across Datasets (mean across trials*±*standard deviation)

### 4.3 Monkey Bump Task with Ephys from Area 2 of Somatosensory Cortex

Single-trial neural dynamics have become increasingly important for understanding how the brain responds to varied behaviorally-relevant events. We applied SSLD to a single-trial electrophysiology dataset collected from a monkey performing a bump task (16 conditions with 8 active and 8 passive) [Chowdhury et al., 2020]. Neural activity was recorded from Brodmann’s Area 2 of somatosensory cortex (*D*_*n*_ = 65), and was aligned to movement onset with a 500*ms* window before and after movement [Pei et al., 2021]. The behavioral observations included hand position, hand velocity, and force applied to the arm (*D*_*b*_ = 10). The dataset contained 193 active trials and 171 passive trials. More details can be found in Appendix B. SSLD performed well on both active and passive trials; here, we mainly report the active-trial results.

Consistent with the reaching dataset, SSLD accurately reconstructed neural activity and behavior (Fig. 3A,B; see Table G.1 for quantitative results). Linear decoding showed that behavioral information was concentrated in the shared and private behavioral latents, but largely absent from the private neural latent (Fig. 3C). Decoding performance using an LSTM decoder is also reported in Table H.1. The shared latent also captured the global task geometry, with its correlation structure closely matching behavioral condition similarity (Fig. 3D and Fig. M.7B) and outperforming neural activity and private latents (Fig. 3E, see Appendix F for details). The inferred discrete states aligned with task-defined behavioral epochs (Fig. 3F, Table 1). Using *D*_*s*_ = 8, the shared latent trajectories showed clear condition-level organization (Fig. 3G), consistent with hand-movement trajectories across trials (Fig. 3H).

**Figure 3:**
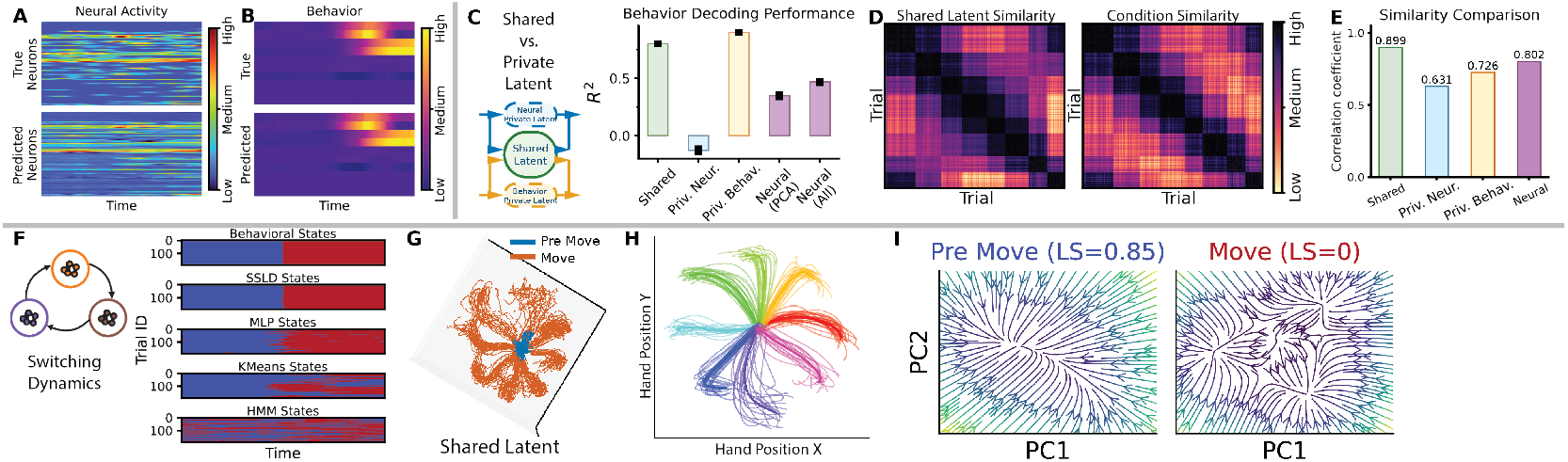
Bump Task: Representative reconstructions of (A) neural activity and (B) behavior. *Shared vs. private latent*: (C) Behavioral information is mainly encoded in the shared and private behavioral latent, as assessed using a linear decoder (see Table H.1 for LSTM decoder). (D) The shared-latent similarity matrix and condition similarity across the 8 active conditions. (E) Linear regression coefficients relating condition similarity to similarity in the shared latent, private neural latent, private behavioral latent, and raw neural activity, respectively (see Appendix F). *Switching dynamics*: (F) Inferred discrete states align with ‘pre move’ and ‘move’. (G) Shared latent trajectories show structured organization across active-trial conditions, consistent with the corresponding (H) hand-movement trajectories. (I) Flow fields of the RNN in distinct behavioral states (LS in the titles), reveal state-dependent latent dynamics.

We also visualized shared-latent flow fields across inferred states for active trials (Fig. 3I) and quantified them using line-attractor scores (see Fig. M.12 for passive trials). The flow fields differed across behavioral epochs and trial types. In active trials, the line-attractor score was higher before movement onset and decreased during movement, consistent with the reaching dataset. Ablation analyses also mirrored the reaching results: CCA generalized poorly to the test set, removing the behavioral encoder and decoder reduced behavioral decoding, and removing switching dynamics slightly lowered state-alignment accuracy (Table 2). These results support the roles of shared latent structure, behavioral supervision, and switching dynamics in recovering neural-behavioral dynamics.

### 4.4 Mouse Self-Initiated Decision-Making Task with WFCI from Dorsal Cortex

As modern recording technologies increasingly capture neural activity across multiple brain regions, there is a growing need to understand how distributed neural dynamics relate to behavior. We next show the performance of SSLD on a self-initiated decision-making task, with single-trial neural activity recorded using WFCI (*D*_*n*_ = 200) across 22 brain regions. The behavioral data contains pose estimation of the paws, nose, and tongue (*D*_*b*_ = 8). The details of this dataset and pre-processing can be found in Appendix of this paper and Musall et al. [2019]. SSLD accurately reconstructed neural activity and behavior (Fig. 4A,B; see Table G.1 for quantitative results), and its inferred discrete states were well aligned with the task-defined behavioral epochs (Fig. 4C, Table 1). Using *D*_*s*_ = 16, the shared latent trajectories showed clear condition-level organization in the learned neural-behavioral representation (Fig. 4D). Behavior decoding showed that behavioral information was concentrated in the shared and private behavioral latents, but largely absent from the private neural latent (Fig. 4E and Table H.1.). We visualized RNN flow fields across inferred states and quantified them using line-attractor scores (Fig. M.13). The score increased after stimulus onset and exceeded 1, suggesting that the stimulus leads to the presence of an accumulation mode in neural dynamics.

**Figure 4:**
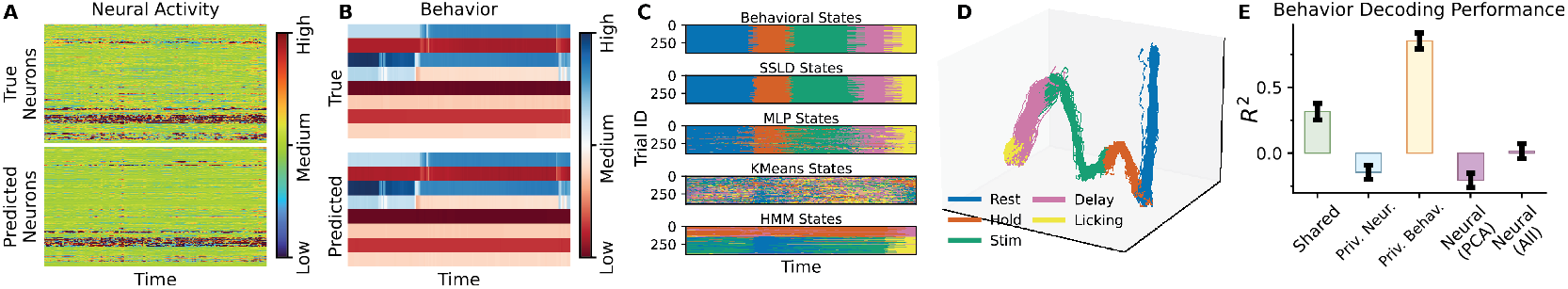
Decision Making: Reconstructions of (A) neural activity and (B) behavior. (C) Inferred discrete states align with behavioral states. (D) Shared latent trajectories. (E) Behavioral information is mainly encoded in the shared and private behavioral latents (see Table H.1 for LSTM decoder).

Finally, ablation analyses showed trends consistent with the previous datasets. A linear CCA-based shared subspace failed to generalize well, removing the behavioral encoder and decoder reduced behavioral decoding performance, and removing the switching recurrent structure led to lower statealignment accuracy (Table 2). These results again highlight the importance of the shared latent structure, behavioral supervision, and switching dynamics.

### 4.5 Mouse Spontaneous Movements with WFCI from Dorsal Cortex

We finally evaluate SSLD on spontaneous movements, which involve more complex discrete state transitions due to the mouse’s unconstrained behavior. Neural activity was recorded using WFCI, details are provided in the Appendix B.

Consistent with the previous datasets, SSLD accurately reconstructed both neural population activity and behavior, as shown by examples in Fig. 5A and Fig. 5B (see Table G.1 for quantitative results). The inferred discrete states show partial alignment with the labeled behavioral epochs (Fig. 5C, Table 1), suggesting that SSLD captures behaviorally structured trends in the neural dynamics, although state recovery is less accurate in this more naturalistic experimental paradigm. For the main analysis, we used a shared latent dimensionality of *D*_*s*_ = 16 and further evaluated the effect of *D*_*s*_ in Fig. M.5. The shared latent trajectories show clear organization across behavioral conditions in the learned neural-behavioral representation (Fig. 5D). We next examined how behavioral information was distributed across the learned representations. As in the previous datasets, behavior decoding showed that behavioral information is primarily represented in the shared latent and private behavioral latent, while remaining largely absent from the private neural latent (Fig. 5E and Table H.1.). We also visualize the RNN flow fields in Fig. M.14 and find that the dynamics remain stable across different spontaneous behavioral states.

**Figure 5:**
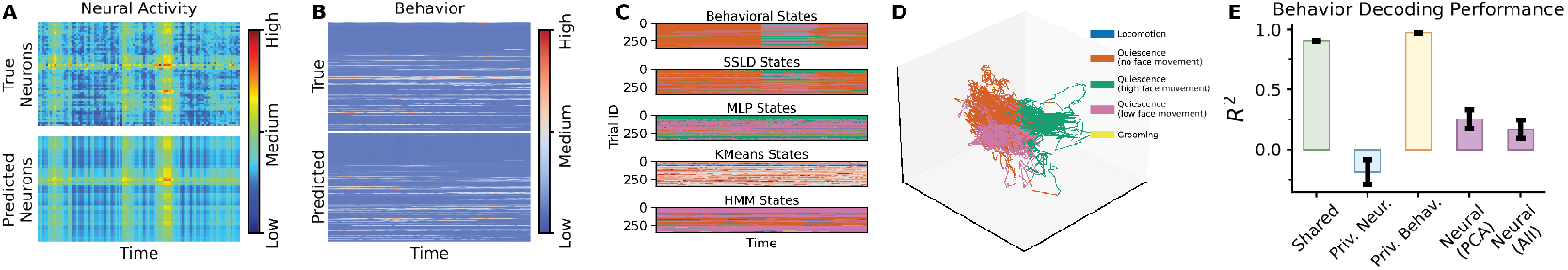
Spontaneous Movement: Representative reconstructions of (A) neural activity and (B) behavior. (C) Inferred discrete states align with task-defined behavioral epochs. (D) Shared latent. (E) Behavioral information is mainly encoded in the shared and private behavioral latents (see Table H.1 for LSTM decoder).

## 5 Conclusion and Future Work

Here, we introduce SSLD, a model that combines shared neural-behavioral representation learning with switching dynamics. By placing switching in the shared latent space, SSLD isolates behaviorally relevant neural dynamics and captures structured state transitions. Across datasets, it performs accurate reconstruction, aligns inferred states with behavior, and yields interpretable latent dynamics. These results suggest that jointly modeling shared structure and switching dynamics is key for linking neural activity to behavior. Several limitations should be noted. First, the model relies on sufficient prior knowledge of behavior, including well-defined discrete states and high-quality behavioral recordings; insufficient or poorly characterized behavioral information may degrade model performance. Second, the current framework treats neural-behavioral coupling as internally driven, without explicitly modeling external inputs or interventions. Future work will address this by incorporating input-driven or causal components to account for external perturbations, and will extend the model to integrate additional modalities such as video for a more comprehensive representation of neural-behavioral relationships. No negative societal impact is noted.

## Acknowledgement

We gratefully acknowledge support from the NIH R34DA059718 and NSF NCS 2350329 to SS, R01MH099045, R01MH113852, and DP1EY033975 to MJH, R01EY022951 to JAC, R01MH113852 to MJH and JAC, and EY026878 to the Yale Vision Core. We also acknowledge support from the Grand Challenge program of the Wu Tsai Institute at Yale University.

## A Simulated Datasets

We randomly generate two dynamical systems and use them to simulate a sequence of 100 time points. The first 50 time points are generated from the first dynamical system, and the remaining 50 time points are generated from the second dynamical system, creating a known switching point in the middle of the sequence. Private neural and behavioral latents are independently sampled from Gaussian noise, while the generated switching latent is treated as the ground-truth shared latent. To generate observations, we concatenate the shared latent with the private neural latent and pass it through a linear readout to obtain simulated neural activity. Similarly, we concatenate the shared latent with the private behavioral latent and pass it through another linear readout to obtain simulated behavior. We randomly generate 50 trials, using 40 trials for training and 10 trials for testing.

## B Experimental Datasets

### Electrophysiology Recordings during a Monkey Reaching Task

In the reaching dataset, the recording starts with the monkey in a resting state, and a target shows up after 400ms. The monkey then gets a ‘go’ cue after a delay. Thereafter, the monkey starts moving from the origin to the targets and hold at the targets upon reaching. More details can be found in Churchland et al. [2012]. We explore the relationship between the change of neural dynamics and the change of behavioral states. Here, we use the trial-average neural activity which is averaged across multiple trials with the same conditions, details of the trial-average data can be found in Churchland et al. [2012]. The data is recorded for 180 neurons and has 236 time points representing 2.36 seconds. We have 5 different behaviorally-relevant states: (1) resting; (2) delay before the ‘go’ cue; (3) reaction time (from receiving ‘go’ cue to the start of movement); (4) movement execution; (5) hold at the target. This dataset is publicly available: https://dandiarchive.org/dandiset/000128

### Electrophysiology Recordings during a Monkey Bump Task

In Area 2 bump dataset, neural activity was recorded from Brodmann’s Area 2 of primary somatosensory cortex in a non-human primate during a center-out reaching task with mechanical perturbations Chowdhury et al. [2020]. In this task, the monkey performed both active reaches and passive movements, where external forces (‘bumps’) were applied to the arm prior to movement initiation. Neural activity was recorded as single-unit spiking and converted to firing rates, while behavioral measurements included hand position, velocity, and applied forces, capturing the kinematics of the entire arm. The dataset contains multiple movement conditions spanning different directions and perturbation types, providing a rich testbed for studying how somatosensory cortex encodes both internally generated and externally driven movements. We analyzed single-trial data across 16 conditions (8 active, 8 passive), aligned to movement onset with a 500*ms* window before and after movement. The number of time points in each trial is 200. We use code provided from [Pei et al., 2021] to process the datasets, the kernal size to smooth the neural spiking data is 40. The behavioral epochs were defined as (1) ‘pre move’ and (2) ‘move’ periods. This dataset is publicly available: https://dandiarchive.org/dandiset/000129/0.241017.1444

### Widefield Calcium Imaging during Mouse Decision-Making Behavior

In self-initiated decision making dataset, mice initiated trials by touching either of two handles and hold a handle for 1*s* followed by sensory stimuli. The sensory stimulus was presented for 600ms, and after a 500ms pause with no stimulus and then the stimulus was repeated for another 600ms. After the second stimulus, a 1000ms delay was imposed, then the mice are required to lick one of two spouts, the mice were rewarded with a drop of water if they licked the spout twice on the same side as the stimulus. More details can be found in Musall et al. [2019]. The data is processed via LocaNMF Saxena et al. [2020], and the 200 temporal components returned by LocaNMF are used. The data has 189 time points representing 6.3 seconds. In this data, mice have 5 behaviorally-relevant states, i.e., (1) from baseline to holding of the handle; (2) holding the handle; (3) stimulus presentation; (4) delay from the end of stimulus to the start of licking; (5) licking the spout. This dataset is publicly available: https://repository.cshl.edu/id/eprint/38599/

### Widefield Calcium Imaging during Mouse Spontaneous Movements

In the spontaneous movements dataset, cortical activity was recorded from awake mice using wide-field mesoscopic calcium imaging during unconstrained spontaneous behavior [Benisty et al., 2024]. The dataset consists of continuous neural recordings rather than stimulus-aligned or trial-based activity. Neural activity was represented using cortical parcels defined from the CCFv3 atlas [Wang et al., 2020], providing region-level calcium activity traces across dorsal cortex. Simultaneously recorded behavioral variables include measures related to facial motion, pupil dynamics, and locomotion. Here, we use the preprocessed parcel-level calcium traces and segment the continuous recordings into temporal samples for model fitting. Because the dataset does not contain discrete trials, we treat each temporal segment as one sample of spontaneous cortical dynamics. More details of the imaging acquisition, behavioral recordings, cortical parcellation, and preprocessing can be found in Benisty et al. [2024]. In this dataset, mice freely run on a cylindrical wheel, and wheel speed is used as the behavioral variable. During spontaneous movement, mice exhibit 5 behaviorally relevant states: i.e., (1) locomotion; (2) quiescence with no face movement; (3) quiescence with high face movement; (4) quiescence with low face movement; (5) grooming. This dataset is publicly available: https://dandiarchive.org/dandiset/001560/0.250829.1249

## C Choice of Shared Latent Dimensionality

We selected the shared latent dimensionality *D*_*s*_ based on four criteria: reconstruction performance for neural activity and behavior, accuracy in recovering behavioral states, and the amount of behavioral information encoded in the private neural latent. Specifically, we chose a value of *D*_*s*_ that achieved strong reconstruction and state-recovery performance while minimizing behavioral decoding performance from the private neural latent. High behavioral information in the private neural latent would indicate incomplete disentanglement between shared and modality-specific representations. SSLD performance across different choices of *D*_*s*_ for the four datasets is shown in Fig. M.2, Fig. M.3, Fig. M.4, and Fig. M.5.

## D Optional Initialization

We provide an optional initialization in which SSLD prioritizes the shared latent to reconstruct both neural activity and behavior. This encourages faster convergence and helps avoid poor local minima, such as solutions where information is predominantly encoded in private latents. We apply this initialization when training takes a long time to converge.

## E Linear Decoding of Behavior from Latent Representations

To evaluate how well different representations capture behaviorally relevant information, we trained linear decoders to predict behavior from several sources. Specifically, we fit linear regression models to decode behavioral variables from the shared latent, private neural and private behavioral latent, raw neural activity, and PCA-reduced neural activity where the number of principal components was matched to the shared latent dimensionality (*D*_*s*_) to ensure a fair comparison.

## F Task Geometry in Shared Latent

In both reaching and bump tasks, we tested whether the shared latent captures the global geometry of the behavioral task. We defined a behavioral condition-similarity matrix using the inverse Euclidean distance between source-to-target reaching paths and compared it with the correlation structure of the inferred shared latents across task conditions. In both datasets, the two structures (Fig. 2D and Fig. 3D) were highly aligned.

We then applied linear regression between behavioral condition similarity and shared latent similarity (Fig. M.6B and Fig. M.7B). As a result, conditions with similar task geometry produced more similar shared-latent representations, reflected by a high correlation coefficient.

Compared with the similarity of raw neural activity, private behavioral latents, and private neural latents (Fig. 2E and Fig. 3E), the shared latent showed the strongest correspondence with behavioral condition similarity, indicating that it preferentially captures global task structure.

## G Neural and Behavioral reconstruction using SSLD

**Table G.1:** Mean squared error (MSE) of neural and behavioral reconstruction on each dataset.

| Reconstruction | Reaching | Bump | Decision-making | Spontaneous |
|---|---|---|---|---|
| Neural | 0.003 ± 0.002 | 0.0022 ± 0.0003 | 0.000095 ± 0.000093 | 0.0003 ± 0.001 |
| Behavioral | 0.002 ± 0.003 | 0.0002 ± 0.0001 | 0.0001 ± 0.0002 | 0.000096 ± 0.000027 |

## H Behavior Decoding from Latent using Nonlinear Decoder

**Table H.1:**
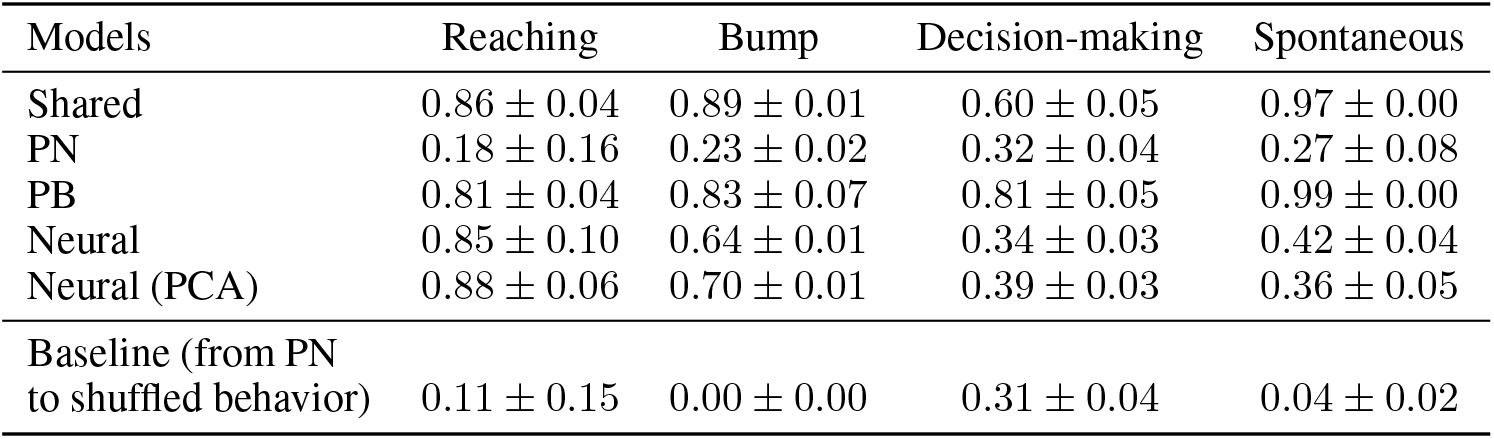
Behavior Decoding using an LSTM Decoder.

## I Comparison with other foundation models

**Table I.1:** Behavior Decoding Performance.

| Datasets<br>Models | Reaching | Bump | Decision-making | Spontaneous |
|---|---|---|---|---|
| POYO | 0.74 ± 0.11 | 0.57 ± 0.02 | 0.11 ± 0.06 | 0.29 ± 1.16 |
| **SSLD (LSTM decoder)** | **0.86 ± 0.04** | **0.89 ± 0.01** | **0.60 ± 0.05** | **0.97 ± 0.00** |

## J Unsupervised SSLD

**Table J.1:** Behavior Decoding using Shared Latent across Datasets.

| Datasets<br>Models | Reaching | Bump | Decision-making | Spontaneous |
|---|---|---|---|---|
| Unsupervised SSLD | 0.36±0.15 | **0.80±0.07** | **0.39±1.46** | 0.86±0.39 |
| **SSLD** | **0.51±0.12** | **0.80±0.09** | 0.32±1.26 | **0.91±0.14** |

## K Comparison between SSLD with different objective function

We compared different disentanglement objectives by evaluating the state accuracy and behavioral decoding performance using an LSTM decoder. We tested private latent with dimensionality either equal to or half that of the shared latent. These settings corresponded to 8 or 4 dimensions for the reaching and bump datasets, and 16 or 8 dimensions for the decision-making and spontaneous movement datasets.

**Table K.1:** State accuracy and Shared latent behavioral decoding performance of SSLD with different disentanglement objectives, evaluated with an LSTM decoder.

| Dataset | Metric | Objective |  |  |  |  |
|---|---|---|---|---|---|---|
|  |  | CS | Orthogonality |  | Total correlation |  |
|  |  |  | Same Dim | Lower Dim | Same Dim | Lower Dim |
| Reaching | State accuracy | 0.96 ± 0.04 | 0.95 ± 0.04 | 0.95 ± 0.06 | **0.96 ± 0.02** | 0.96 ± 0.03 |
|  | Behavior decoding | **0.86 ± 0.04** | 0.80 ± 0.12 | 0.81 ± 0.11 | 0.84 ± 0.07 | 0.84 ± 0.09 |
| Bump | State accuracy | **0.99 ± 0.00** | **0.99 ± 0.00** | **0.99 ± 0.00** | **0.99 ± 0.00** | **0.99 ± 0.00** |
|  | Behavior decoding | **0.89 ± 0.01** | **0.89 ± 0.01** | 0.88 ± 0.22 | 0.82 ± 0.06 | 0.87 ± 0.02 |
| Decision-making | State accuracy | 0.95 ± 0.04 | 0.95 ± 0.05 | **0.96 ± 0.04** | 0.91 ± 0.06 | 0.91 ± 0.05 |
|  | Behavior decoding | 0.60 ± 0.05 | **0.62 ± 0.06** | 0.57 ± 0.05 | 0.60 ± 0.05 | 0.36 ± 0.04 |
| Spontaneous | State accuracy | **0.51 ± 0.23** | 0.51 ± 0.24 | 0.48 ± 0.26 | 0.47 ± 0.24 | 0.47 ± 0.23 |
|  | Behavior decoding | 0.97 ± 0.00 | 0.57 ± 0.05 | 0.96 ± 0.00 | **0.98 ± 0.00** | **0.98 ± 0.00** |

## L Details of SSLD Structure

**Table L.1:**
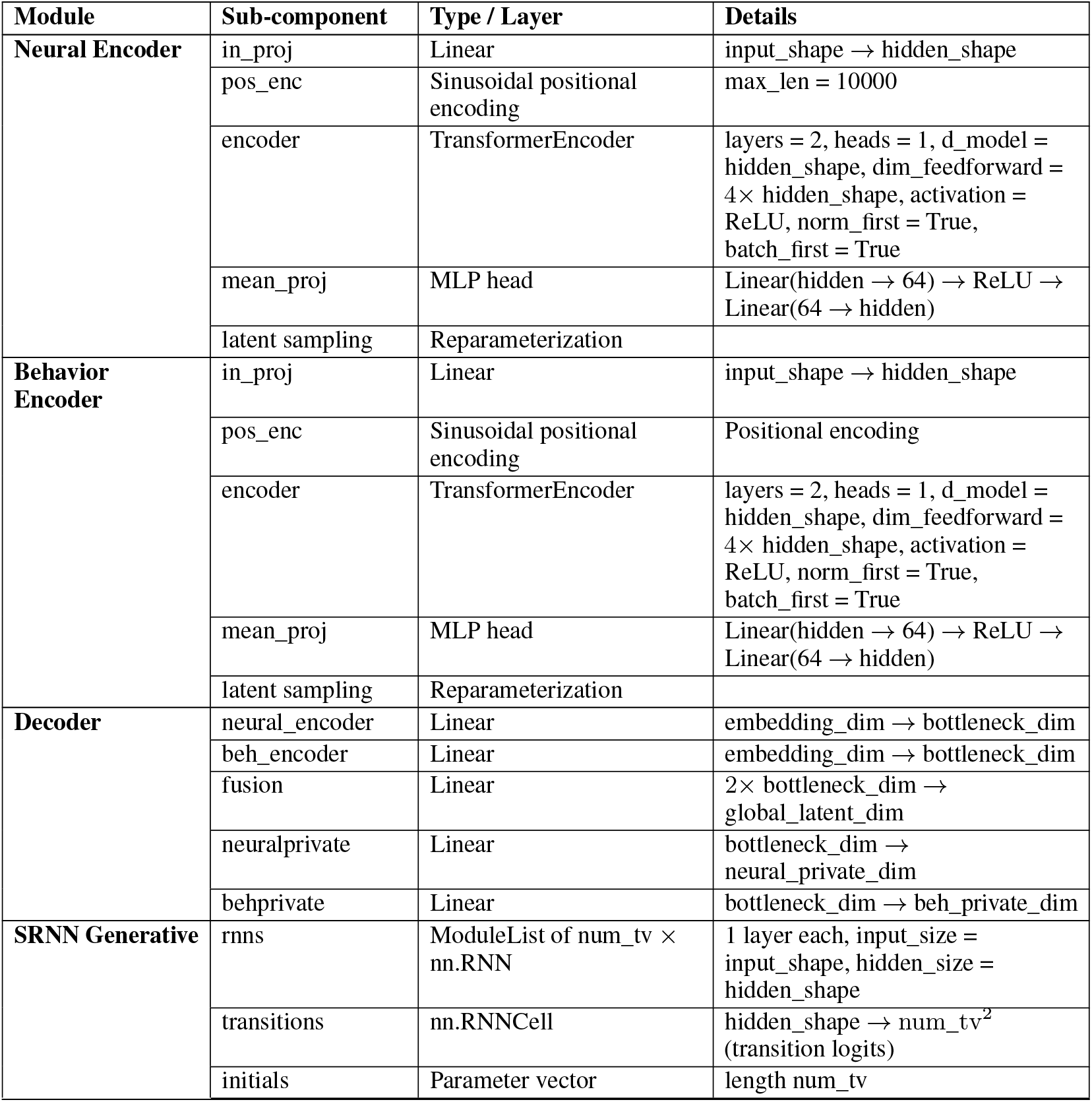
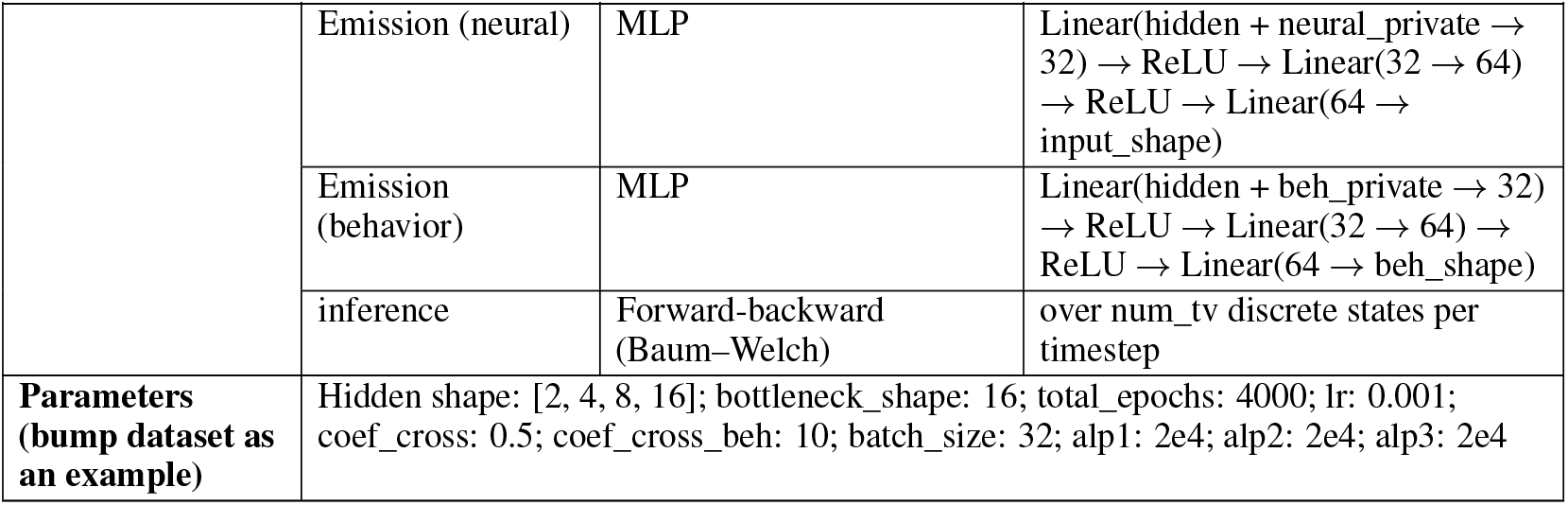
Model architecture and training parameters.

| Module | Sub-component | Type / Layer | Details |
|---|---|---|---|
| **Neural Encoder** | in_proj | Linear | input_shape → hidden_shape |
|  | pos_enc | Sinusoidal positional encoding | max_len = 10000 |
|  | encoder | TransformerEncoder | layers = 2, heads = 1, d_model = hidden_shape, dim_feedforward = 4× hidden_shape, activation = ReLU, norm_first = True, batch_first = True |
|  | mean_proj | MLP head | Linear(hidden → 64) → ReLU → Linear(64 → hidden) |
|  | latent sampling | Reparameterization |  |
| **Behavior Encoder** | in_proj | Linear | input_shape → hidden_shape |
|  | pos_enc | Sinusoidal positional encoding | Positional encoding |
|  | encoder | TransformerEncoder | layers = 2, heads = 1, d_model = hidden_shape, dim_feedforward = 4× hidden_shape, activation = ReLU, norm_first = True, batch_first = True |
|  | mean_proj | MLP head | Linear(hidden → 64) → ReLU → Linear(64 → hidden) |
|  | latent sampling | Reparameterization |  |
| **Decoder** | neural_encoder | Linear | embedding_dim → bottleneck_dim |
|  | beh_encoder | Linear | embedding_dim → bottleneck_dim |
|  | fusion | Linear | 2× bottleneck_dim → global_latent_dim |
|  | neuralprivate | Linear | bottleneck_dim → neural_private_dim |
|  | behprivate | Linear | bottleneck_dim → beh_private_dim |
| **SRNN Generative** | rnns | ModuleList of num_tv × nn.RNN | 1 layer each, input_size = input_shape, hidden_size = hidden_shape |
|  | transitions | nn.RNNCell | hidden_shape → num_tv<sup>2</sup> (transition logits) |
|  | initials | Parameter vector | length num_tv |
|  | Emission (neural) | MLP | Linear(hidden + neural_private → 32) → ReLU → Linear(32 → 64) → ReLU → Linear(64 → input_shape) |
|  | Emission (behavior) | MLP | Linear(hidden + beh_private → 32) → ReLU → Linear(32 → 64) → ReLU → Linear(64 → beh_shape) |
|  | inference | Forward-backward (Baum–Welch) | over num_tv discrete states per timestep |
| **Parameters (bump dataset as an example)** | Hidden shape: [2, 4, 8, 16]; bottleneck_shape: 16; total_epochs: 4000; lr: 0.001; coef_cross: 0.5; coef_cross_beh: 10; batch_size: 32; alp1: 2e4; alp2: 2e4; alp3: 2e4 |  |  |

## M Appendix Figures

**Figure M.1:**
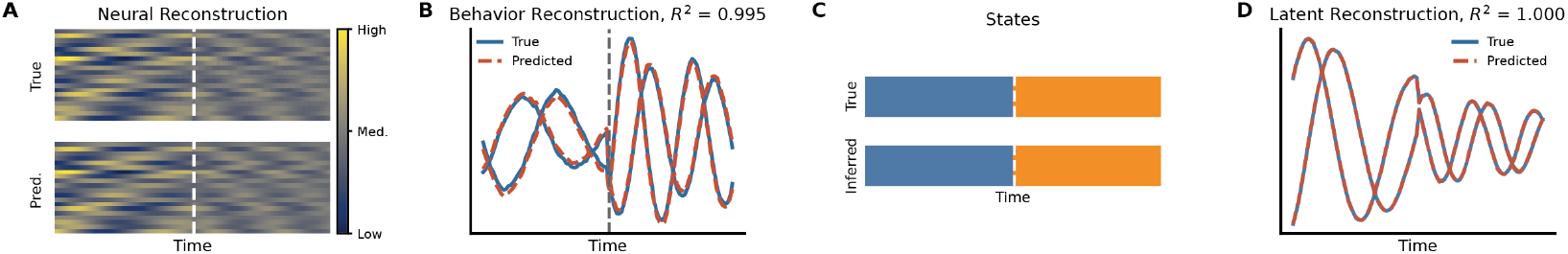
Simulated: (A) Reconstruction of simulated activity and (B) behavior using SSLD. (C) Comparison between ground truth discrete states and SSLD-recovered discrete states. (D) Comparison between ground truth latent and SSLD-recovered shared latent (after linear transformation).

**Figure M.2:**
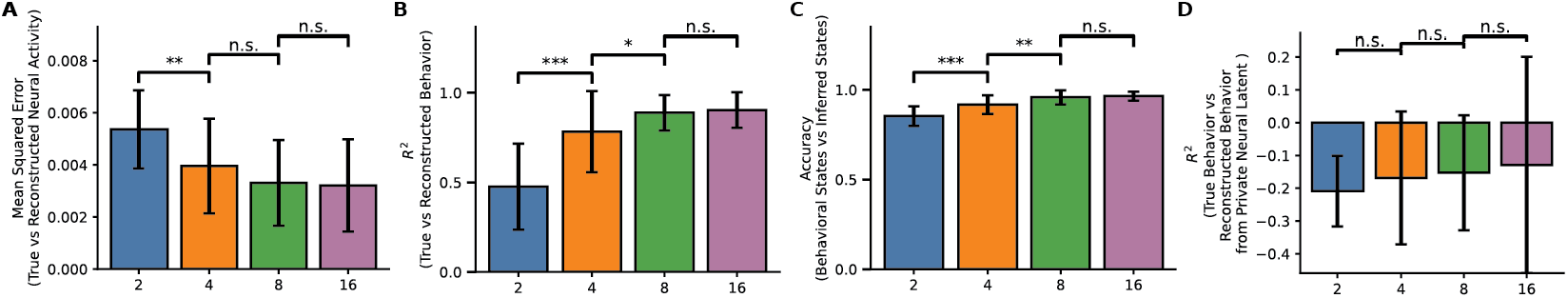
Reaching: Model performance as a function of shared latent dimensionality *D*_*s*_. (A) Neural activity reconstruction. (B) Behavioral reconstruction. (C) Behavioral state recovery accuracy. (D) Behavioral decoding from the private neural latent representation.

**Figure M.3:**
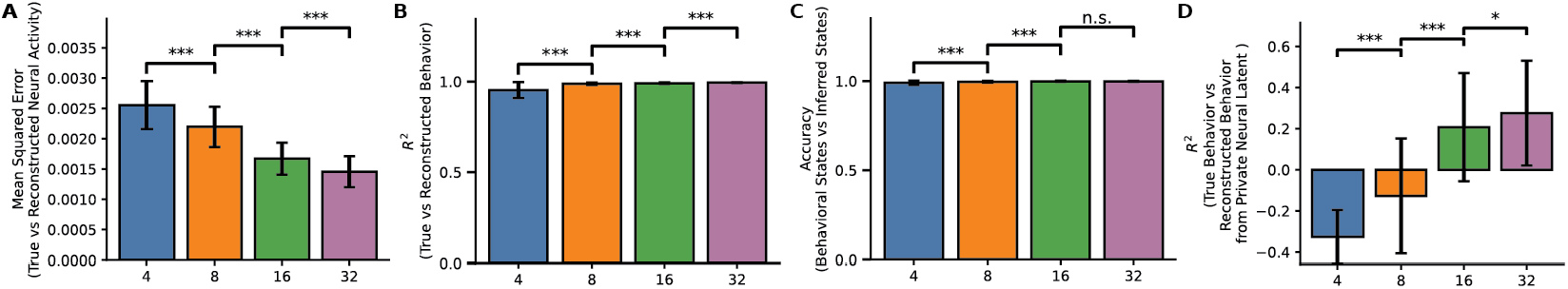
Bump Task: Model performance as a function of shared latent dimensionality *D*_*s*_. (A) Neural activity reconstruction. (B) Behavioral reconstruction. (C) Behavioral state recovery accuracy. (D) Behavioral decoding from the private neural latent representation.

**Figure M.4:**
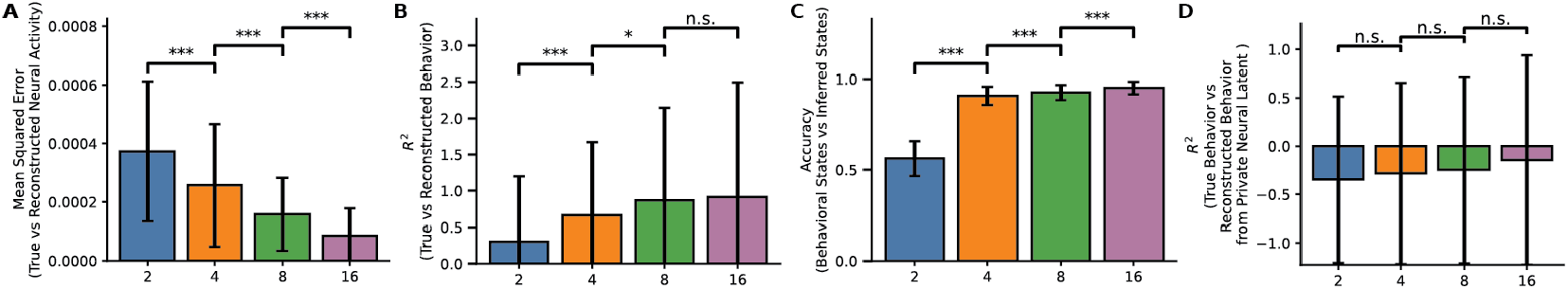
Decision Making: Model performance as a function of shared latent dimensionality *D*_*s*_. (A) Neural activity reconstruction. (B) Behavioral reconstruction. (C) Behavioral state recovery accuracy. (D) Behavioral decoding from the private neural latent representation.

**Figure M.5:**
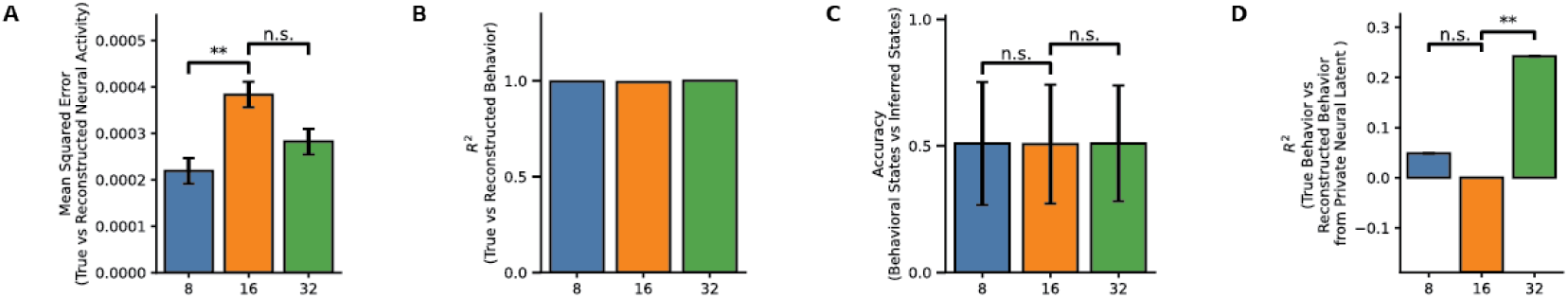
Spontaneous Movement: Model performance as a function of shared latent dimensionality *D*_*s*_. (A) Neural activity reconstruction. (B) Behavioral reconstruction. (C) Behavioral state recovery accuracy. (D) Behavioral decoding from the private neural latent representation.

**Figure M.6:**
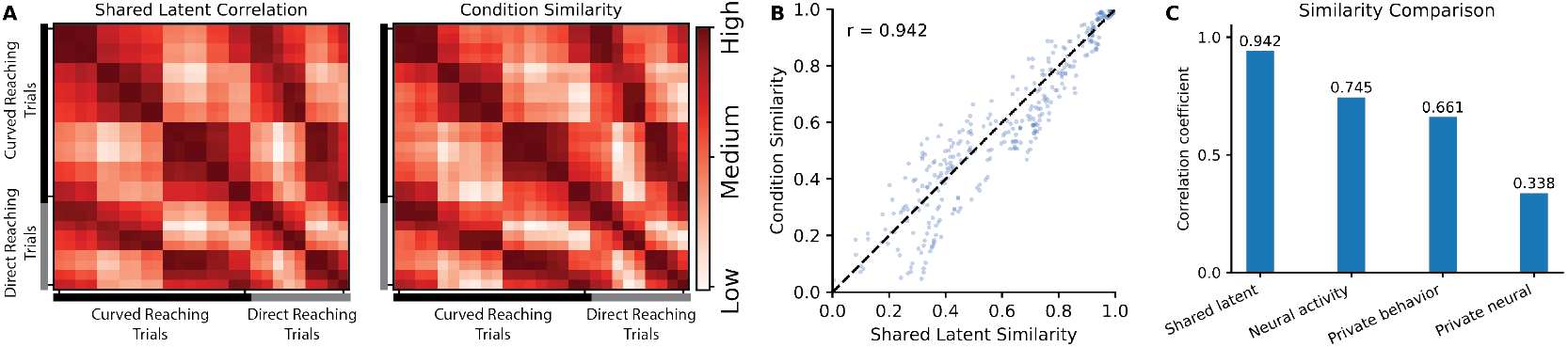
Reaching: (A) The shared-latent correlation matrix and condition similarity across the 27 reaching conditions. Behavioral condition-similarity matrix is defined using the inverse Euclidean distance between source-to-target reaching paths. (B) Linear regression between the two matrices in (A). (C) Linear regression coefficient for raw neural activity, the private behavioral latent, and the private neural latent. The shared latent showed the strongest correspondence with behavioral condition similarity, indicating that it preferentially captures global task structure.

**Figure M.7:**
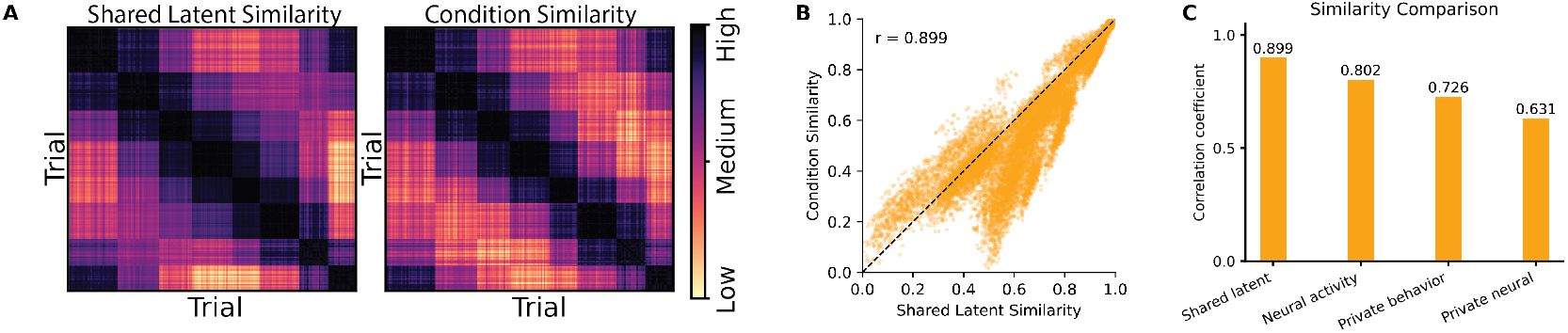
Bump Task: (A) The shared-latent correlation matrix and condition similarity across the 8 active conditions. (B) Linear regression between the two matrices in (A). (C) Linear regression coefficient for raw neural activity, the private behavioral latent, and the private neural latent.

**Figure M.8:**
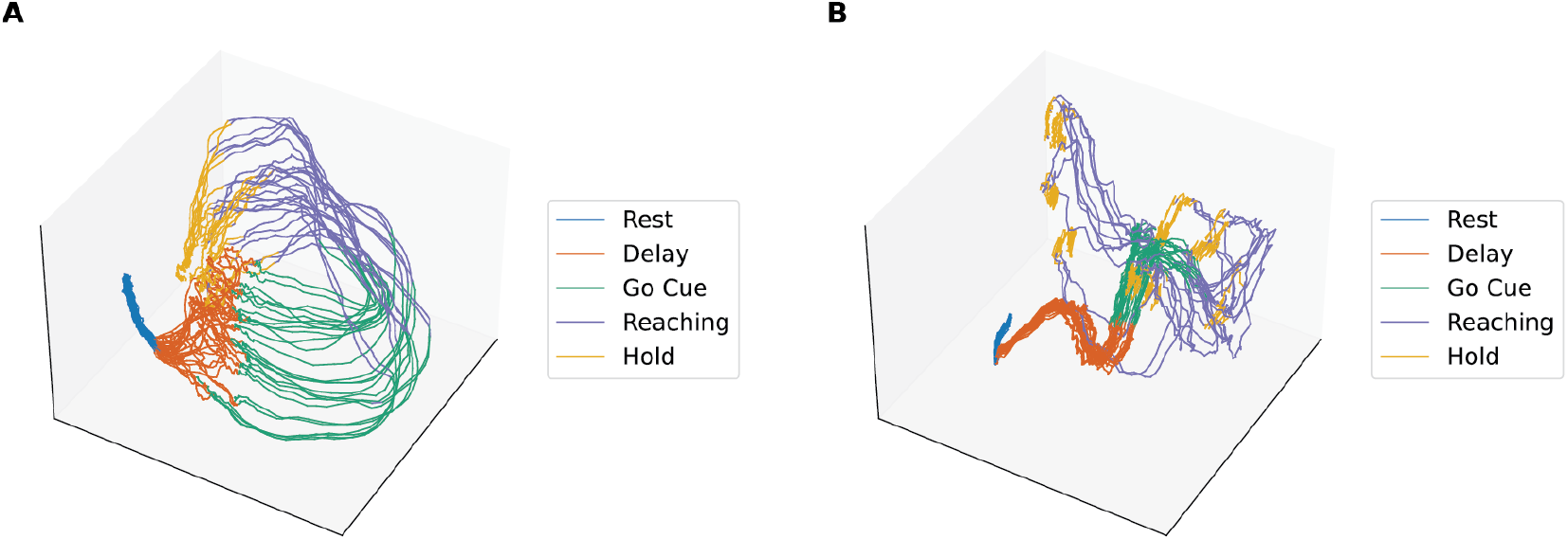
Reaching: Visualization of the learned (A) private neural latent and (B) private behavioral latent representations, colored according to behavioral states.

**Figure M.9:**
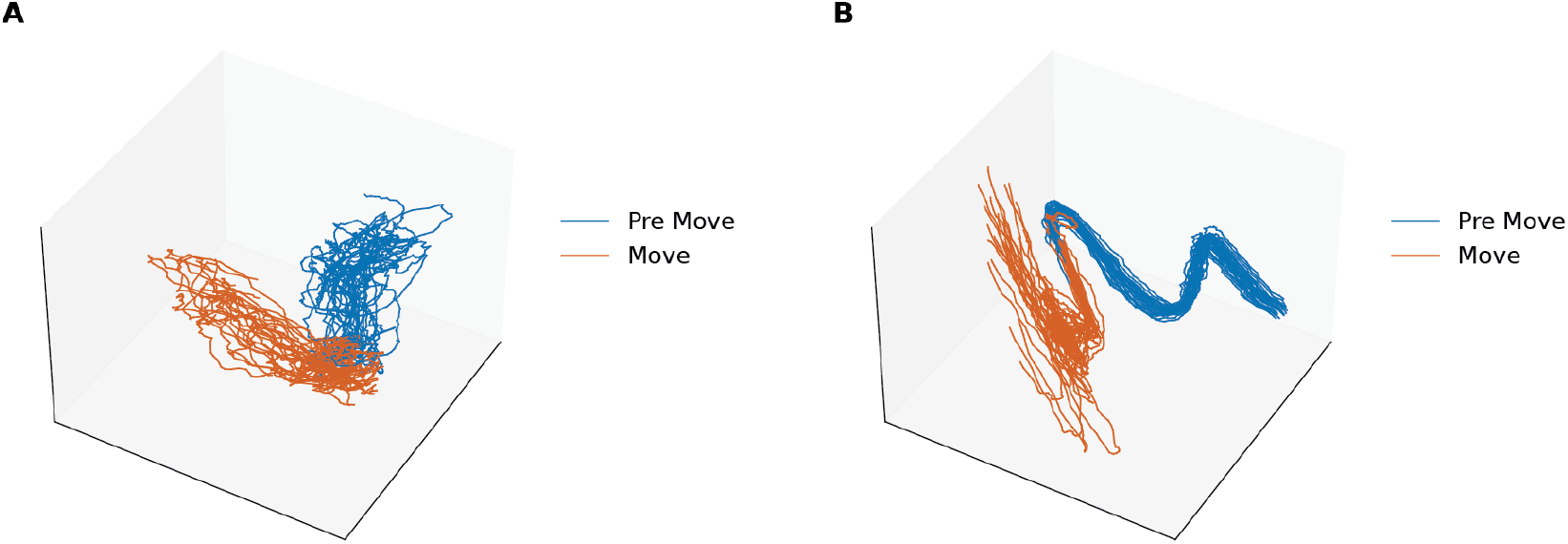
Bump Task: Visualization of the learned (A) private neural latent and (B) private behavioral latent representations.

**Figure M.10:**
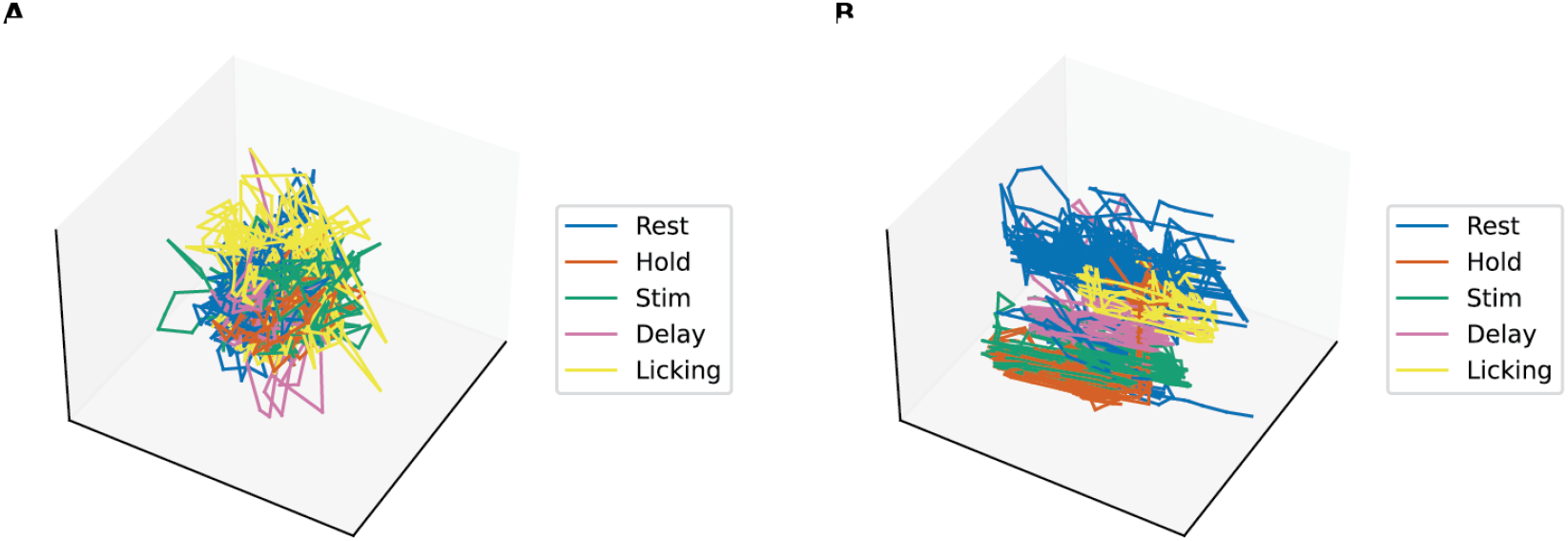
Decision Making: Visualization of the learned (A) private neural latent and (B) private behavioral latent representations.

**Figure M.11:**
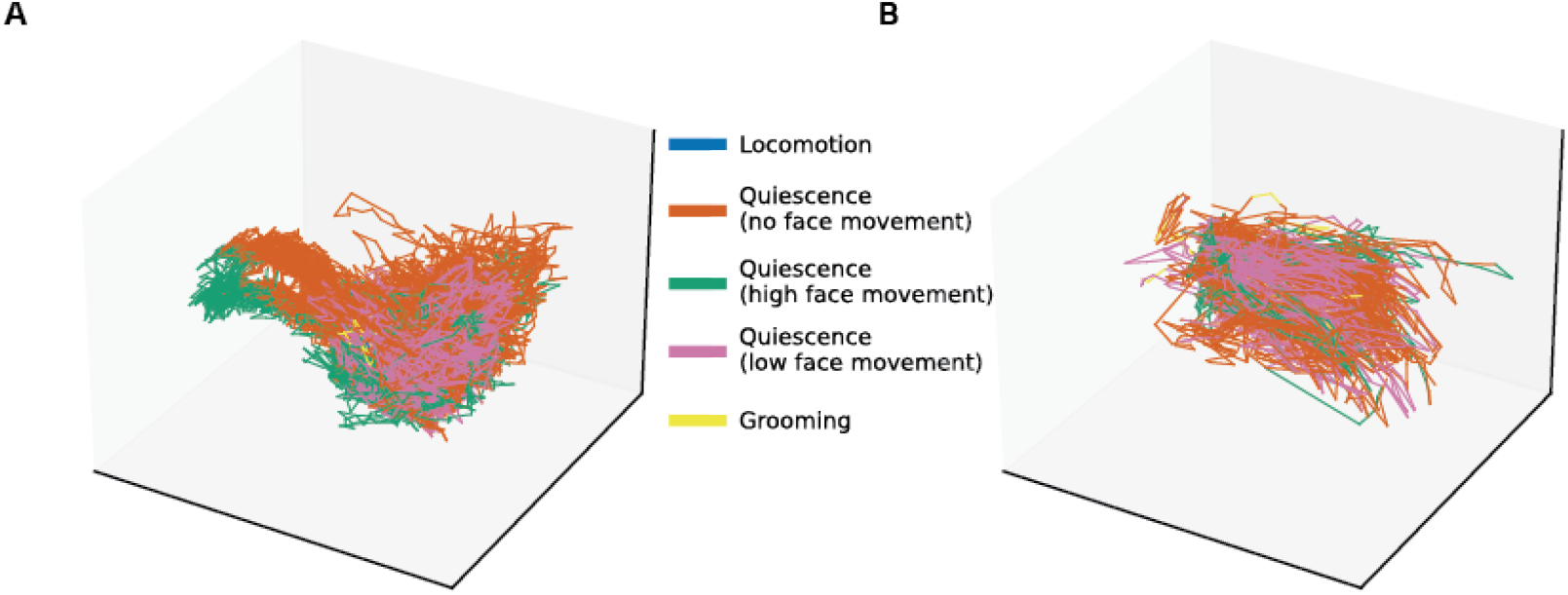
Spontaneous Movement: Visualization of the learned (A) private neural latent and (B) private behavioral latent representations.

**Figure M.12:**
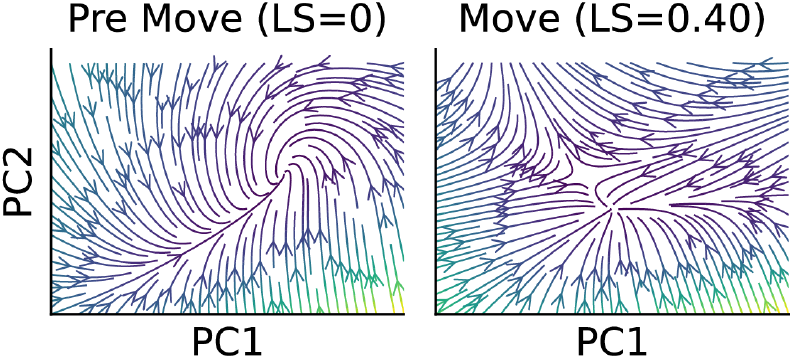
Bump Task: Flow fields of RNNs for passive trials. The passive trials show distinct dynamics from active trials, likely due to the external perturbation.

**Figure M.13:**
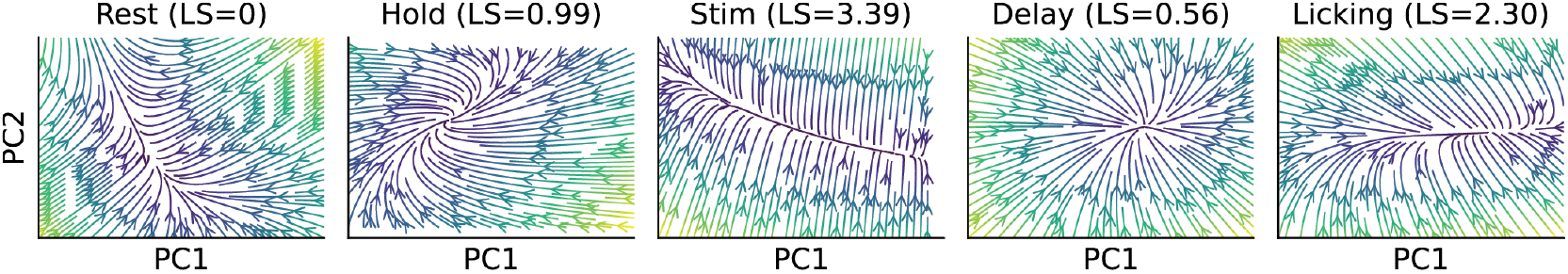
Decision Making: Flow fields of RNNs

**Figure M.14:**
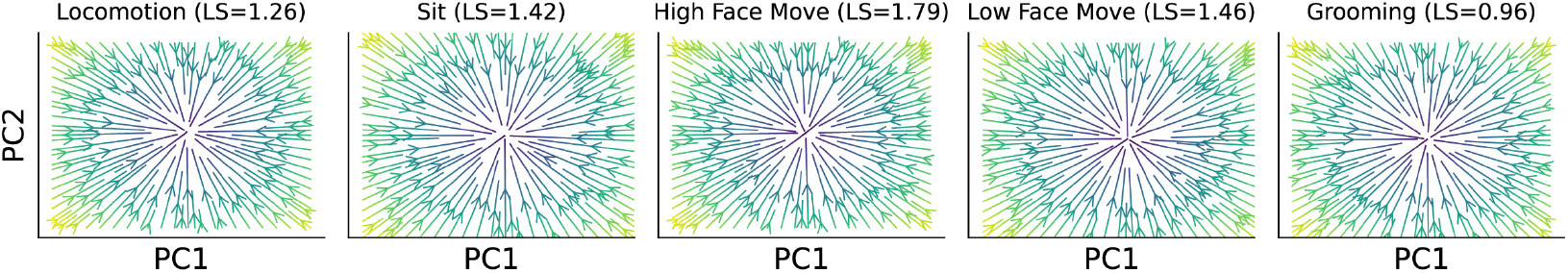
Spontaneous Movement: Flow fields of RNNs

## NeurIPS Paper Checklist

The checklist is designed to encourage best practices for responsible machine learning research, addressing issues of reproducibility, transparency, research ethics, and societal impact. Do not remove the checklist: **The papers not including the checklist will be desk rejected**. The checklist should follow the references and follow the (optional) supplemental material. The checklist does NOT count towards the page limit.

Please read the checklist guidelines carefully for information on how to answer these questions. For each question in the checklist:

- You should answer [Yes], [No], or [N/A].
- [N/A] means either that the question is Not Applicable for that particular paper or the relevant information is Not Available.
- Please provide a short (1–2 sentence) justification right after your answer (even for [N/A]).

### The checklist answers are an integral part of your paper submission

They are visible to the reviewers, area chairs, senior area chairs, and ethics reviewers. You will also be asked to include it (after eventual revisions) with the final version of your paper, and its final version will be published with the paper.

The reviewers of your paper will be asked to use the checklist as one of the factors in their evaluation. While [Yes] is generally preferable to [No], it is perfectly acceptable to answer [No] provided a proper justification is given (e.g., error bars are not reported because it would be too computationally expensive” or “we were unable to find the license for the dataset we used”). In general, answering [No] or [N/A] is not grounds for rejection. While the questions are phrased in a binary way, we acknowledge that the true answer is often more nuanced, so please just use your best judgment and write a justification to elaborate. All supporting evidence can appear either in the main paper or the supplemental material, provided in appendix. If you answer [Yes] to a question, in the justification please point to the section(s) where related material for the question can be found.

1. **Claims** Question: Do the main claims made in the abstract and introduction accurately reflect the paper’s contributions and scope? Answer: [Yes] Justification: The abstract and introduction accurately reflect the paper’s contributions, we focus on the core methodological advance of Switching Shared Latent Dynamics (SSLD): combining shared neural–behavioral representation learning with nonlinear switching dynamics. We evaluated the model on simulated and experimental datasets, the model can successfully recover behaviorally aligned states, separate shared and private latent structure, and ablation analyses support the model components. Guidelines:
  - The answer [N/A] means that the abstract and introduction do not include the claims made in the paper.
  - The abstract and/or introduction should clearly state the claims made, including the contributions made in the paper and important assumptions and limitations. A [No] or [N/A] answer to this question will not be perceived well by the reviewers.
  - The claims made should match theoretical and experimental results, and reflect how much the results can be expected to generalize to other settings.
  - It is fine to include aspirational goals as motivation as long as it is clear that these goals are not attained by the paper.
2. **Limitations** Question: Does the paper discuss the limitations of the work performed by the authors? Answer: [Yes] Justification: We discuss two limitations in Section of conclusion and future works. Guidelines:
  - The answer [N/A] means that the paper has no limitation while the answer [No] means that the paper has limitations, but those are not discussed in the paper.
  - The authors are encouraged to create a separate “Limitations” section in their paper.
  - The paper should point out any strong assumptions and how robust the results are to violations of these assumptions (e.g., independence assumptions, noiseless settings, model well-specification, asymptotic approximations only holding locally). The authors should reflect on how these assumptions might be violated in practice and what the implications would be.
  - The authors should reflect on the scope of the claims made, e.g., if the approach was only tested on a few datasets or with a few runs. In general, empirical results often depend on implicit assumptions, which should be articulated.
  - The authors should reflect on the factors that influence the performance of the approach. For example, a facial recognition algorithm may perform poorly when image resolution is low or images are taken in low lighting. Or a speech-to-text system might not be used reliably to provide closed captions for online lectures because it fails to handle technical jargon.
  - The authors should discuss the computational efficiency of the proposed algorithms and how they scale with dataset size.
  - If applicable, the authors should discuss possible limitations of their approach to address problems of privacy and fairness.
  - While the authors might fear that complete honesty about limitations might be used by reviewers as grounds for rejection, a worse outcome might be that reviewers discover limitations that aren’t acknowledged in the paper. The authors should use their best judgment and recognize that individual actions in favor of transparency play an important role in developing norms that preserve the integrity of the community. Reviewers will be specifically instructed to not penalize honesty concerning limitations.
3. **Theory assumptions and proofs** Question: For each theoretical result, does the paper provide the full set of assumptions and a complete (and correct) proof? Answer: [Yes] Justification: We provide equations on the main text for our model and training. Guidelines:
  - The answer [N/A] means that the paper does not include theoretical results.
  - All the theorems, formulas, and proofs in the paper should be numbered and crossreferenced.
  - All assumptions should be clearly stated or referenced in the statement of any theorems.
  - The proofs can either appear in the main paper or the supplemental material, but if they appear in the supplemental material, the authors are encouraged to provide a short proof sketch to provide intuition.
  - Inversely, any informal proof provided in the core of the paper should be complemented by formal proofs provided in appendix or supplemental material.
  - Theorems and Lemmas that the proof relies upon should be properly referenced.
4. **Experimental result reproducibility** Question: Does the paper fully disclose all the information needed to reproduce the main experimental results of the paper to the extent that it affects the main claims and/or conclusions of the paper (regardless of whether the code and data are provided or not)? Answer: [Yes] Justification: We cite all the source of our datasets in the main text and include the details of our experiments in Method and Result section, and we will provide our code for reproduction upon publication. Guidelines:
  - The answer [N/A] means that the paper does not include experiments.
  - If the paper includes experiments, a [No] answer to this question will not be perceived well by the reviewers: Making the paper reproducible is important, regardless of whether the code and data are provided or not.
  - If the contribution is a dataset and/or model, the authors should describe the steps taken to make their results reproducible or verifiable.
  - Depending on the contribution, reproducibility can be accomplished in various ways. For example, if the contribution is a novel architecture, describing the architecture fully might suffice, or if the contribution is a specific model and empirical evaluation, it may be necessary to either make it possible for others to replicate the model with the same dataset, or provide access to the model. In general. releasing code and data is often one good way to accomplish this, but reproducibility can also be provided via detailed instructions for how to replicate the results, access to a hosted model (e.g., in the case of a large language model), releasing of a model checkpoint, or other means that are appropriate to the research performed.
  - While NeurIPS does not require releasing code, the conference does require all submissions to provide some reasonable avenue for reproducibility, which may depend on the nature of the contribution. For example
    a. If the contribution is primarily a new algorithm, the paper should make it clear how to reproduce that algorithm.
    b. If the contribution is primarily a new model architecture, the paper should describe the architecture clearly and fully.
    c. If the contribution is a new model (e.g., a large language model), then there should either be a way to access this model for reproducing the results or a way to reproduce the model (e.g., with an open-source dataset or instructions for how to construct the dataset).
    d. We recognize that reproducibility may be tricky in some cases, in which case authors are welcome to describe the particular way they provide for reproducibility. In the case of closed-source models, it may be that access to the model is limited in some way (e.g., to registered users), but it should be possible for other researchers to have some path to reproducing or verifying the results.
5. **Open access to data and code** Question: Does the paper provide open access to the data and code, with sufficient instructions to faithfully reproduce the main experimental results, as described in supplemental material? Answer: [Yes] Justification: We cite all the source of our datasets which are all in the public domain; we provide our full code for reproduction in the paper. Guidelines:
  - The answer [N/A] means that paper does not include experiments requiring code.
  - Please see the NeurIPS code and data submission guidelines (https://neurips.cc/public/guides/CodeSubmissionPolicy) for more details.
  - While we encourage the release of code and data, we understand that this might not be possible, so [No] is an acceptable answer. Papers cannot be rejected simply for not including code, unless this is central to the contribution (e.g., for a new open-source benchmark).
  - The instructions should contain the exact command and environment needed to run to reproduce the results. See the NeurIPS code and data submission guidelines (https://neurips.cc/public/guides/CodeSubmissionPolicy) for more details.
  - The authors should provide instructions on data access and preparation, including how to access the raw data, preprocessed data, intermediate data, and generated data, etc.
  - The authors should provide scripts to reproduce all experimental results for the new proposed method and baselines. If only a subset of experiments are reproducible, they should state which ones are omitted from the script and why.
  - At submission time, to preserve anonymity, the authors should release anonymized versions (if applicable).
  - Providing as much information as possible in supplemental material (appended to the paper) is recommended, but including URLs to data and code is permitted.
6. **Experimental setting/details** Question: Does the paper specify all the training and test details (e.g., data splits, hyperpa-rameters, how they were chosen, type of optimizer) necessary to understand the results? Answer: [Yes] Justification: We provide details in the Methods and Results section. Guidelines:
  - The answer [N/A] means that the paper does not include experiments.
  - The experimental setting should be presented in the core of the paper to a level of detail that is necessary to appreciate the results and make sense of them.
  - The full details can be provided either with the code, in appendix, or as supplemental material.
7. **Experiment statistical significance** Question: Does the paper report error bars suitably and correctly defined or other appropriate information about the statistical significance of the experiments? Answer: [Yes] Justification: We provide details in the Results section. Guidelines:
  - The answer [N/A] means that the paper does not include experiments.
  - The authors should answer [Yes] if the results are accompanied by error bars, confidence intervals, or statistical significance tests, at least for the experiments that support the main claims of the paper.
  - The factors of variability that the error bars are capturing should be clearly stated (for example, train/test split, initialization, random drawing of some parameter, or overall run with given experimental conditions).
  - The method for calculating the error bars should be explained (closed form formula, call to a library function, bootstrap, etc.)
  - The assumptions made should be given (e.g., Normally distributed errors).
  - It should be clear whether the error bar is the standard deviation or the standard error of the mean.
  - It is OK to report 1-sigma error bars, but one should state it. The authors should preferably report a 2-sigma error bar than state that they have a 96% CI, if the hypothesis of Normality of errors is not verified.
  - For asymmetric distributions, the authors should be careful not to show in tables or figures symmetric error bars that would yield results that are out of range (e.g., negative error rates).
  - If error bars are reported in tables or plots, the authors should explain in the text how they were calculated and reference the corresponding figures or tables in the text.
8. **Experiments compute resources** Question: For each experiment, does the paper provide sufficient information on the computer resources (type of compute workers, memory, time of execution) needed to reproduce the experiments? Answer: [Yes] Justification: We provide details in the Methods section. Guidelines:
  - The answer [N/A] means that the paper does not include experiments.
  - The paper should indicate the type of compute workers CPU or GPU, internal cluster, or cloud provider, including relevant memory and storage.
  - The paper should provide the amount of compute required for each of the individual experimental runs as well as estimate the total compute.
  - The paper should disclose whether the full research project required more compute than the experiments reported in the paper (e.g., preliminary or failed experiments that didn’t make it into the paper).
9. **Code of ethics** Question: Does the research conducted in the paper conform, in every respect, with the NeurIPS Code of Ethics https://neurips.cc/public/EthicsGuidelines? Answer: [Yes] Justification: Our research conforms with the NeurIPS Code of Ethics. Guidelines:
  - The answer [N/A] means that the authors have not reviewed the NeurIPS Code of Ethics.
  - If the authors answer [No], they should explain the special circumstances that require a deviation from the Code of Ethics.
  - The authors should make sure to preserve anonymity (e.g., if there is a special consideration due to laws or regulations in their jurisdiction).
10. **Broader impacts** Question: Does the paper discuss both potential positive societal impacts and negative societal impacts of the work performed? Answer: [Yes] Justification: This is addressed in the Conclusions section. Guidelines:
  - The answer [N/A] means that there is no societal impact of the work performed.
  - If the authors answer [N/A] or [No], they should explain why their work has no societal impact or why the paper does not address societal impact.
  - Examples of negative societal impacts include potential malicious or unintended uses (e.g., disinformation, generating fake profiles, surveillance), fairness considerations (e.g., deployment of technologies that could make decisions that unfairly impact specific groups), privacy considerations, and security considerations.
  - The conference expects that many papers will be foundational research and not tied to particular applications, let alone deployments. However, if there is a direct path to any negative applications, the authors should point it out. For example, it is legitimate to point out that an improvement in the quality of generative models could be used to generate Deepfakes for disinformation. On the other hand, it is not needed to point out that a generic algorithm for optimizing neural networks could enable people to train models that generate Deepfakes faster.
  - The authors should consider possible harms that could arise when the technology is being used as intended and functioning correctly, harms that could arise when the technology is being used as intended but gives incorrect results, and harms following from (intentional or unintentional) misuse of the technology.
  - If there are negative societal impacts, the authors could also discuss possible mitigation strategies (e.g., gated release of models, providing defenses in addition to attacks, mechanisms for monitoring misuse, mechanisms to monitor how a system learns from feedback over time, improving the efficiency and accessibility of ML).
11. **Safeguards** Question: Does the paper describe safeguards that have been put in place for responsible release of data or models that have a high risk for misuse (e.g., pre-trained language models, image generators, or scraped datasets)? Answer: [N/A] Justification: The paper poses no such risks. Guidelines:
  - The answer [N/A] means that the paper poses no such risks.
  - Released models that have a high risk for misuse or dual-use should be released with necessary safeguards to allow for controlled use of the model, for example by requiring that users adhere to usage guidelines or restrictions to access the model or implementing safety filters.
  - Datasets that have been scraped from the Internet could pose safety risks. The authors should describe how they avoided releasing unsafe images.
  - We recognize that providing effective safeguards is challenging, and many papers do not require this, but we encourage authors to take this into account and make a best faith effort.
12. **Licenses for existing assets** Question: Are the creators or original owners of assets (e.g., code, data, models), used in the paper, properly credited and are the license and terms of use explicitly mentioned and properly respected? Answer: [Yes] Justification: We have cited all the source of our datasets. Guidelines:
  - The answer [N/A] means that the paper does not use existing assets.
  - The authors should cite the original paper that produced the code package or dataset.
  - The authors should state which version of the asset is used and, if possible, include a URL.
  - The name of the license (e.g., CC-BY 4.0) should be included for each asset.
  - For scraped data from a particular source (e.g., website), the copyright and terms of service of that source should be provided.
  - If assets are released, the license, copyright information, and terms of use in the package should be provided. For popular datasets, paperswithcode.com/datasets has curated licenses for some datasets. Their licensing guide can help determine the license of a dataset.
  - For existing datasets that are re-packaged, both the original license and the license of the derived asset (if it has changed) should be provided.
  - If this information is not available online, the authors are encouraged to reach out to the asset’s creators.
13. **New assets** Question: Are new assets introduced in the paper well documented and is the documentation provided alongside the assets? Answer: [N/A] Justification: The paper does not release new assets. Guidelines:
  - The answer [N/A] means that the paper does not release new assets.
  - Researchers should communicate the details of the dataset/code/model as part of their submissions via structured templates. This includes details about training, license, limitations, etc.
  - The paper should discuss whether and how consent was obtained from people whose asset is used.
  - At submission time, remember to anonymize your assets (if applicable). You can either create an anonymized URL or include an anonymized zip file.
14. **Crowdsourcing and research with human subjects** Question: For crowdsourcing experiments and research with human subjects, does the paper include the full text of instructions given to participants and screenshots, if applicable, as well as details about compensation (if any)? Answer: [N/A] Justification: The paper does not involve crowdsourcing nor research with human subjects Guidelines:
  - The answer [N/A] means that the paper does not involve crowdsourcing nor research with human subjects.
  - Including this information in the supplemental material is fine, but if the main contribution of the paper involves human subjects, then as much detail as possible should be included in the main paper.
  - According to the NeurIPS Code of Ethics, workers involved in data collection, curation, or other labor should be paid at least the minimum wage in the country of the data collector.
15. **Institutional review board (IRB) approvals or equivalent for research with human subjects** Question: Does the paper describe potential risks incurred by study participants, whether such risks were disclosed to the subjects, and whether Institutional Review Board (IRB) approvals (or an equivalent approval/review based on the requirements of your country or institution) were obtained? Answer: [N/A] Justification: The paper does not involve crowdsourcing nor research with human subjects. Guidelines:
  - The answer [N/A] means that the paper does not involve crowdsourcing nor research with human subjects.
  - Depending on the country in which research is conducted, IRB approval (or equivalent) may be required for any human subjects research. If you obtained IRB approval, you should clearly state this in the paper.
  - We recognize that the procedures for this may vary significantly between institutions and locations, and we expect authors to adhere to the NeurIPS Code of Ethics and the guidelines for their institution.
  - For initial submissions, do not include any information that would break anonymity (if applicable), such as the institution conducting the review.
16. **Declaration of LLM usage** Question: Does the paper describe the usage of LLMs if it is an important, original, or non-standard component of the core methods in this research? Note that if the LLM is used only for writing, editing, or formatting purposes and does *not* impact the core methodology, scientific rigor, or originality of the research, declaration is not required. Answer: [N/A] Justification: The paper does not involve LLMs as any important, original, or non-standard components. Guidelines:
  - The answer [N/A] means that the core method development in this research does not involve LLMs as any important, original, or non-standard components.
  - Please refer to our LLM policy in the NeurIPS handbook for what should or should not be described.

## References

M. Azabou, V. Arora, V. Ganesh, X. Mao, S. Nachimuthu, M. Mendelson, B. Richards, M. Perich, G. Lajoie, and E. Dyer. A unified, scalable framework for neural population decoding. Advances in Neural Information Processing Systems, 36:44937–44956, 2023.

M. Azabou, K. X. Pan, V. Arora, I. J. Knight, E. L. Dyer, and B. A. Richards. Multi-session, multi-task neural decoding from distinct cell-types and brain regions. In The Thirteenth International Conference on Learning Representations, 2025.

H. Benisty, D. Barson, A. H. Moberly, S. Lohani, L. Tang, R. R. Coifman, M. C. Crair, G. Mishne, J. A. Cardin, and M. J. Higley. Rapid fluctuations in functional connectivity of cortical networks encode spontaneous behavior. Nature Neuroscience, 27(1):148–158, 2024. doi: 10.1038/s41593-023-01498-y.

K. Bousmalis, G. Trigeorgis, N. Silberman, D. Krishnan, and D. Erhan. Domain separation networks. Advances in neural information processing systems, 29, 2016.

J. O. Caro, A. H. de Oliveira Fonseca, S. A. Rizvi, M. Rosati, C. Averill, J. L. Cross, P. Mittal, E. Zappala, R. M. Dhodapkar, C. Abdallah, et al. Brainlm: A foundation model for brain activity recordings. In The Twelfth International Conference on Learning Representations, 2024.

G. Chau, C. Wang, S. Talukder, V. Subramaniam, S. Soedarmadji, Y. Yue, B. Katz, and A. Barbu. Population transformer: Learning population-level representations of neural activity. ArXiv, pages arXiv–2406, 2025.

R. H. Chowdhury, J. I. Glaser, and L. E. Miller. Area 2 of primary somatosensory cortex encodes kinematics of the whole arm. Elife, 9:e48198, 2020.

M. M. Churchland, J. P. Cunningham, M. T. Kaufman, J. D. Foster, P. Nuyujukian, S. I. Ryu, and K. V. Shenoy. Neural population dynamics during reaching. Nature, 487(7405):51–56, 2012.

Z. Dong, B. Seybold, K. Murphy, and H. Bui. Collapsed amortized variational inference for switching nonlinear dynamical systems. In International Conference on Machine Learning, pages 2638–2647. PMLR, 2020.

D. Durstewitz, G. Koppe, and M. I. Thurm. Reconstructing computational system dynamics from neural data with recurrent neural networks. Nature Reviews Neuroscience, 24(11):693–710, 2023.

J. Glaser, M. Whiteway, J. P. Cunningham, L. Paninski, and S. Linderman. Recurrent switching dynamical systems models for multiple interacting neural populations. Advances in neural information processing systems, 33:14867–14878, 2020.

R. Gondur, U. B. Sikandar, E. Schaffer, M. C. Aoi, and S. L. Keeley. Multi-modal gaussian process variational autoencoders for neural and behavioral data. In The Twelfth International Conference on Learning Representations, 2024.

O. Karniol-Tambour, D. Zoltowski, E. M. Diamanti, L. Pinto, C. Brody, D. Tank, and J. Pillow. Modeling state-dependent communication between brain regions with switching nonlinear dynamical systems. In International Conference on Learning Representations, volume 2024, pages 27010–27030, 2024.

M. T. Kaufman, J. S. Seely, D. Sussillo, S. I. Ryu, K. V. Shenoy, and M. M. Churchland. The largest response component in the motor cortex reflects movement timing but not movement type. eneuro, 3(4), 2016.

D. P. Kingma and M. Welling. Auto-encoding variational bayes. arXiv preprint arXiv:1312.6114, 2013.

M. Lee and V. Pavlovic. Private-shared disentangled multimodal vae for learning of hybrid latent representations. arXiv preprint arXiv:2012.13024, 2020.

S. Linderman, M. Johnson, A. Miller, R. Adams, D. Blei, and L. Paninski. Bayesian learning and inference in recurrent switching linear dynamical systems. In Artificial intelligence and statistics, pages 914–922. PMLR, 2017.

N. Maheswaranathan, A. Williams, M. Golub, S. Ganguli, and D. Sussillo. Reverse engineering recurrent networks for sentiment classification reveals line attractor dynamics. Advances in neural information processing systems, 32, 2019.

S. Musall, M. T. Kaufman, A. L. Juavinett, S. Gluf, and A. K. Churchland. Single-trial neural dynamics are dominated by richly varied movements. Nature neuroscience, 22(10):1677–1686, 2019.

C. Pandarinath, D. J. O’Shea, J. Collins, R. Jozefowicz, S. D. Stavisky, J. C. Kao, E. M. Trautmann, M. T. Kaufman, S. I. Ryu, L. R. Hochberg, et al. Inferring single-trial neural population dynamics using sequential auto-encoders. Nature methods, 15(10):805–815, 2018.

F. Pei, J. Ye, D. Zoltowski, A. Wu, R. H. Chowdhury, H. Sohn, J. E. O’Doherty, K. V. Shenoy, M. T. Kaufman, M. Churchland, et al. Neural latents benchmark’21: evaluating latent variable models of neural population activity. arXiv preprint arXiv:2109.04463, 2021.

A. H.-W. Ryoo, N. H. Krishna, X. Mao, M. Azabou, E. L. Dyer, M. G. Perich, and G. Lajoie. Generalizable, real-time neural decoding with hybrid state-space models. arXiv preprint arXiv:2506.05320, 2025.

O. G. Sani, H. Abbaspourazad, Y. T. Wong, B. Pesaran, and M. M. Shanechi. Modeling behaviorally relevant neural dynamics enabled by preferential subspace identification. Nature Neuroscience, 24 (1):140–149, 2021.

O. G. Sani, B. Pesaran, and M. M. Shanechi. Dissociative and prioritized modeling of behaviorally relevant neural dynamics using recurrent neural networks. Nature neuroscience, 27(10):2033–2045, 2024.

E. Santana, M. Emigh, and J. C. Principe. Information theoretic-learning auto-encoder. In 2016 international joint conference on neural networks (IJCNN), pages 3296–3301. IEEE, 2016.

S. Saxena, I. Kinsella, S. Musall, S. H. Kim, J. Meszaros, D. N. Thibodeaux, C. Kim, J. Cunningham, E. M. Hillman, A. Churchland, et al. Localized semi-nonnegative matrix factorization (locanmf) of widefield calcium imaging data. PLoS computational biology, 16(4):e1007791, 2020.

S. Schneider, J. H. Lee, and M. W. Mathis. Learnable latent embeddings for joint behavioural and neural analysis. Nature, 617(7960):360–368, 2023.

A. Vinograd, A. Nair, J. H. Kim, S. W. Linderman, and D. J. Anderson. Causal evidence of a line attractor encoding an affective state. Nature, 634(8035):910–918, 2024.

Q. Wang, S.-L. Ding, Y. Li, J. Royall, D. Feng, P. Lesnar, N. Graddis, M. Naeemi, B. Facer, A. Ho, et al. The allen mouse brain common coordinate framework: A 3d reference atlas. Cell, 181(4): 936–953.e20, 2020. doi: 10.1016/j.cell.2020.04.007.

J. Ye, J. Collinger, L. Wehbe, and R. Gaunt. Neural data transformer 2: multi-context pretraining for neural spiking activity. Advances in Neural Information Processing Systems, 36:80352–80374, 2023.

D. Yi, H. Dong, M. J. Higley, A. Churchland, and S. Saxena. Shared-ae: Automatic identification of shared subspaces in high-dimensional neural and behavioral activity. In The Thirteenth International Conference on Learning Representations, 2025.

Y. Zhang and S. Saxena. Inference of neural dynamics using switching recurrent neural networks. In The Thirty-eighth Annual Conference on Neural Information Processing Systems, 2024.

Y. Zhang, Y. Wang, D. M. Jiménez-Benetó, Z. Wang, M. Azabou, B. Richards, R. Tung, O. Winter, E. Dyer, L. Paninski, et al. Towards a” universal translator” for neural dynamics at single-cell, single-spike resolution. Advances in Neural Information Processing Systems, 37:80495–80521, 2024.

Y. Zhang, Y. Wang, M. Azabou, A. Andre, Z. Wang, H. Lyu, T. I. B. Laboratory, E. Dyer, L. Paninski, and C. Hurwitz. Neural encoding and decoding at scale. ArXiv,pages arXiv–2504, 2025.

